# Estimating the mean in the space of ranked phylogenetic trees

**DOI:** 10.1101/2023.05.08.539790

**Authors:** Lars Berling, Lena Collienne, Alex Gavryushkin

**Affiliations:** Biological Data Science Lab, School of Mathematics and Statistics, University of Canterbury, Christchurch, New Zealand

## Abstract

Reconstructing evolutionary histories of biological entities, such as genes, cells, organisms, populations, and species, from phenotypic and molecular sequencing data is central to many biological, palaeontological, and biomedical disciplines. Typically, due to uncertainties and incompleteness in data, the true evolutionary history (phylogeny) is challenging to estimate. Statistical modelling approaches address this problem by introducing and studying probability distributions over all possible evolutionary histories. In practice, computational methods are deployed to learn those distributions typically by sampling them. This approach, however, is fundamentally challenging as it requires designing and implementing various statistical methods over a space of phylogenetic trees (or treespace).

Although the problem of developing statistics over a treespace has received substantial attention in the literature and numerous breakthroughs have been made, it remains largely unsolved. The challenge of solving this problem is two-fold: a treespace has non-trivial often counter-intuitive geometry implying that much of classical Euclidean statistics does not immediately apply; many parametrisations of treespace with promising statistical properties are computationally hard, so they cannot be used in data analyses. As a result, there is no single conventional method for estimating even the most fundamental statistics over any treespace, such as mean and variance, and various heuristics are used in practice. Despite the existence of numerous tree summary methods to approximate means of probability distributions over a treespace based on its geometry, and the theoretical promise of this idea, none of the attempts resulted in a practical method for summarising tree samples.

In this paper we present such a method along with useful properties of our chosen treespace while focusing on its impact on phylogenetic analyses of real datasets. We perform an extensive benchmark study and demonstrate that our method outperforms currently most popular methods with respect to a number of important “quality” statistics. Further, we apply our method to three real datasets ranging from cancer evolution to linguistics and find novel insights into corresponding evolutionary problems in all of them. We hence conclude that this treespace is a promising candidate to serve as a foundation for developing statistics over phylogenetic trees analytically, as well as new computational tools for evolutionary data analyses.

## 1. Introduction

Reconstructing evolutionary (or phylogenetic) trees from sequence data is an important task across a variety of areas including evolutionary biology, epidemiology, developmental biology, palaeontology, and linguistics. For this task, there exist four main tree inference paradigms that are based on different analytical approaches [1]: sequence (dis)similarity, parsimony, maximum likelihood, and Bayesian. Distance (or sequence dissimilarity) based tree reconstruction approaches, including the very popular Neighbour Joining algorithm [2], infer trees from a distance matrix between sequences (typically multiple sequence alignments obtained from DNA, RNA, or proteins). Parsimony based approaches are aimed at finding the tree that minimises the number of changes along its branches given sequence data [3, 4] at the tips. Maximum likelihood based tree reconstruction approaches are sampling the likelihood surface over all possible trees with the aim to find the tree (and branch lengths) that maximises the likelihood given sequence data and an evolutionary model [5–7]. Bayesian inference approaches are aimed at sampling (usually using Markov Chain Monte Carlo algorithms) the joint posterior probability distribution of parameters of interest, typically including the tree parameter [8–11].

It is typical for sequence data, on which phylogenetic inference is performed, to contain stochastic noise, errors, and/or omissions. Such uncertainties in data imply uncertainties in the results of a tree inference method, which are quantified using, for example, bootstrap values in the maximum likelihood or posterior support in the Bayesian framework. Furthermore, inference methods can be inconclusive and result in multiple equally “optimal” trees. Examples of these include equally parsimonious trees [12], trees with indistinguishably similar likelihoods [13], and phylogenetic terraces [14, 15].

To overcome these issues in a statistically sound way, Holmes *et al.* [16] initiated a research programme aimed at developing a mathematical framework for statistical analyses over the space of phylogenetic trees. The programme has since received significant attention in the literature with a number of promising treespaces introduced [17–27] and a range of statistical methods developed [28–38]. However, the ultimate milestone of the original programme of introducing fundamental statistics, such as the mean, variance, confidence intervals, for probability distributions over any treespace in a way that would enable a wide range of practical applications remains illusive [23, 39–41].

One idea that cuts through the many successes so far was that of using geometric properties of treespace to introduce statistical notions. Indeed, this idea was developed by Billera *et al.* [18] for treespaces, where means of probability distributions over the treespace have been defined as geometric means, that is, as trees that minimise the sum of squared distances. Matsen [42] extended this idea to shapes. The definition of variance in this case follows naturally, and this approach has been developed in a number of treespaces [30–32, 40, 43, 44]. Geometric ideas [30, 32, 45] enabled an algorithm based on the idea of Sturm [46] to estimate the mean of a set of trees. Nye [29] developed a construction of the first principle component, a milestone for statistical inference in treespaces, with its tropical version developed in [37, 38]. In addition, Barden *et al.* [47] were able to prove a Central Limit Theorem in the BHV [18] treespace with 4 taxa and Nye [33] constructed random walks over this treespace and presented approaches to define uncertainty for collections of phylogenetic trees [35, 36].

These results exposed the main complications that need to be overcome to enable statistically sound and practically useful tree inference methods. These include the stickiness of the mean [41] (irresponsiveness of the mean tree to changes in the tree sample), prevalence of star-like mean trees even for phylogenetically informative data [21, 41], stickiness of the first principal component [25, 29], the high dimensionality problem for small samples of trees [23], and others [34]. Some of these complications are actively being addressed with, e.g. the Wald space [27, 48] being proposed to tackle the mean stickiness issue. These, however, still do not address the problem in full and as of today it is still common practice to use heuristic methods that focus on finding consensus among a given set of trees [40, 49–51]. Partly, this is due to the fact that in treespaces where geometric summary trees were hoped to be phylogenetically informative, such as Robinson-Foulds [43, 52], NNI [53], and SPR [54], they are NP-hard to compute [39, 55, 56] and have hardly received adoption into practice, despite the availability of practical algorithms [57, 58] for small trees and/or distances. In addition, recent results suggest counter-intuitive behaviour of these popularly used metrics [59, 60].

Motivated by the research programme initiated by Holmes *et al.* [16], Gavryushkin & Drummond [21] investigated whether the treespaces developed thus far were readily adaptable to phylogenetic trees scaled to time, which is an important class called time trees with many popular inference methods, such as UPGMA [61], BEAST [9], and BEAST2 [10]. Surprisingly, they discovered that the space of time trees has unique characteristics and is significantly different, geometrically and algorithmically, from the space of classical phylogenetic trees. Along this line of research, it has recently been established that a discrete component of time trees, the RNNI space [24], which is based on tree rearrangement operations, is computationally tractable [62], unlike its classic version the NNI space [55]. Furthermore, it was shown that the RNNI space possesses several characteristics indicating its suitability for statistical approaches [26]. For example, this space satisfies the cluster property [26], which requires a cluster shared by two trees to be also shared by all trees on shortest paths between the two trees. This property is important because it implies that information shared by trees in a sample is preserved in the summary tree. Our results here will justify this further.

In this paper, we continue this line of research and contribute to the programme by demonstrating that for time trees it is possible to introduce fundamental statistics in a way that resolves the main problems (outlined above) to which all known approaches are prone. We demonstrate that for time trees the notion of a mean can be introduced and computationally approximated in a way that is statistically sound, practically computable for large datasets, and an improvement over known approaches commonly used in practice. We carry out a simulation study to validate our method and compare it to other tree summary approaches such as the Maximum Clade Credibility (MCC) method. As part of this study, we have developed an approach to assess suitability of a treespace for statistical analyses, e.g. its “smoothness” with respect to probability distributions over trees. Finally, we apply our method to three real data sets from previous studies of Dravidian languages (linguistics), Weevils (phylogeography), and cancer development (within-organism somatic evolution). Interestingly, in all three cases the summary tree obtained using our methods, although different from the ones published, is consistent with the corresponding evolutionary processes and discussions in the original studies. Combined, these considerations imply that the approach introduced in this paper is a promising candidate to become a mathematical foundation for statistical analyses in the space of phylogenetic time trees.

## 2. Methods

### 2.1. RNNI space

The RNNI space [24] is a treespace of ranked phylogenetic trees, which are rooted binary trees where internal nodes are ordered according to times of the corresponding evolutionary events, assuming no co-occurrence. In other words, every internal node in a ranked tree with *n* leaves is assigned a unique integer (rank) between 1 and *n −* 1 such that no node has rank higher than its parent, and all leaves are assumed to have rank 0. The RNNI space is then defined as a graph where vertices are ranked trees and edges are representing either a rank or an NNI move that transforms one tree into another. A rank move swaps the order of two unconnected nodes with consecutive ranks (i.e. ranks *i* and *i* + 1) in a tree and an NNI move can be performed on an edge in a tree that connects two nodes with consecutive ranks by moving either of the two sister clades adjacent to the bottom node to the opposite side of the top node. The RNNI distance between trees is then defined as the length of a shortest path in this graph. This distance is efficiently computable [62], which enables computational tools necessary for our approaches to be applicable in practice. The RNNI treespace is the main tool used in this paper.

### 2.2. Mean tree

In the following we present an algorithm to approximate the mean (defined as a centroid) tree for a set of ranked trees in the RNNI space. The work of Sturm [46] gives an iterative algorithm to approximate the mean and variance of a probability distribution over a non-positively curved space, which includes treespaces such as the BHV [30, 32, 50], *τ* -space, and *t*-space [21]. They also show that the Law of Large Numbers holds in these spaces, which means that for the sample mean converges to the true mean as the sample size increases. In this paper, we follow the general idea and introduce the Centroid algorithm (Algorithm 1) that uses local tree search to minimise the sum of squared (SoS) distances between a summary tree and a given tree sample and stops when it finds a locally optimal tree *T_a_*, approximating a centroid tree *T ^∗^*.More formally, the input to the Centroid algorithm (Algorithm 1) is a set of trees *T* and a starting tree *T_start_*, which can generally be chosen arbitrarily but is computed by an adaptation of a known tree summary method in our case (see Section 2.2.1 and Supplement for a discussion on this topic). In the first step, *T_a_*:= *T_start_* is used as approximation of the mean tree and the SoS value for *T_start_*is computed. The algorithm proceeds iteratively by computing the SoS values for all neighbours of the current tree *T_a_* (i.e. trees with RNNI distance one to *T_a_*) – this step can be executed in parallel for each neighbour. Then *T_a_* gets updated to be the neighbour with lowest SoS value (if there are multiple such neighbours, one is chosen randomly) and we proceed to the next iteration. If there is no neighbour with lower SoS value than *T_a_*, a local optimum is reached and *T_a_* is returned as approximation of the mean tree for the set *T*.

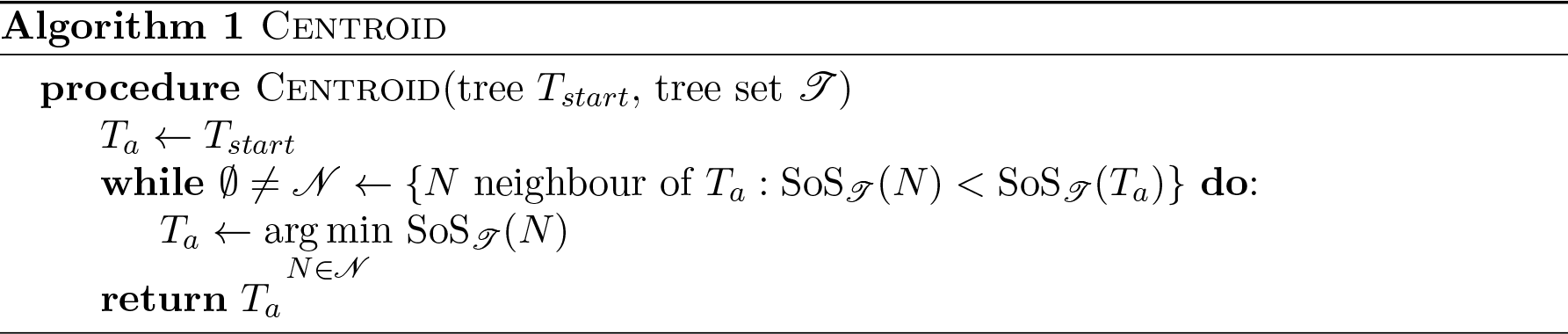

#### 2.2.1. Avoiding non global optima

The Centroid algorithm (Algorithm 1) requires a starting tree as input. Choosing a good initial tree is important for both the accuracy of the algorithm (the quality of the approximation) and the running time. For computing a starting tree for the Centroid algorithm we use the algorithmic idea presented by Sturm [46], which is used in [32] to approximate mean trees in BHV-space and in [21] to approximate the mean in *τ* -space. It was shown by Sturm [46] that this algorithm converges to the true mean under the Law of Large Numbers, and as we’ll see below it provides a good initial starting tree for our approximation algorithm (see Supplement Section 5.1 for a comparison to other choices).

Our version of the Sturm algorithm adapted to the RNNI treespace works as follows. In the first iteration we select a tree *T* uniformly at random from the set of trees *T* and remove it from this set. In every following iteration *k* = 2*, . . ., m* we choose a tree *R* uniformly at random and remove it from *T* . We then compute a shortest path *p* from *R* to *T*, using the FindPath algorithm [62]. The tree *T* for the next iteration is then defined as the tree at position *l* <u>^1^</u> *J* of *p*. Note that this implies that if *l* <u>^1^</u> *J* = 0, the tree *T* does not change. The time complexity of this procedure is in *O*(*|T | ∗ n*^2^) where *O*(*n*^2^) is the complexity of calculating the necessary tree on the shortest path between the two trees in RNNI [62].

#### 2.2.2. Annotating the centroid tree with branch lengths

The approximate mean tree returned by our Centroid algorithm is a ranked tree and hence does not have branch lengths, only an ordering of internal nodes. Most applications, however, require time trees with branch lengths representing evolutionary event times. Because commonly used annotation methods, e.g. from the *TreeAnnotator* programme [9], change the ranking of a tree when annotating it, we cannot use them for our RNNI centroid approximation. We therefore introduce a new annotation method that does not change ranks of internal nodes in the summary tree. Instead of annotating branch lengths based on average heights of the respective clade (set of taxa below the node) as it is done by *TreeAnnotator*, we annotate with the average height of each rank, that is, to calculate the height of the node of rank *i* we take the average of *i*’th *t*-coordinate [21] across the tree sample. For non-ultrametric trees, we extend the *t*-space by extending leaf branches to create an ultrametric tree (similarly to [24]). This approach is a heuristic and we discuss its implications later in the paper (Section 4).

### 2.3. The MCC summary tree heuristic

The *Maximum Clade Credibility* (MCC) tree is computed by the commonly used BEAST [9] utility programme *TreeAnnotator* [63] in a two-step procedure. First the tree topology maximising the posterior clade probability is chosen from the tree sample of the posterior distribution. This implies that an MCC tree contains the set of most probable clades from the sample of trees that is given. This tree is then annotated with clade ages, using one of four different methods. In this paper we consider the *common ancestor* annotation [40]. The other three annotation techniques are discussed in the Supplement (Section 5.8).

### 2.4. Simulation study

To assess the performance of our approach we perform the following study. First, we simulate posterior samples using BEAST2 [10] by generating an initial “true” tree, simulating an alignment down this tree, and then running BEAST2 to compute a posterior sample. We generate a total of 504 different tree sets (i.e. posterior samples) for trees on a range of 50 up to 200 taxa. For each number *n* of taxa we simulate approximately 50 different alignments and corresponding posterior samples, fewer for *n ≥* 100, as the run time of BEAST2 and our Centroid algorithm increases significantly with increasing number of taxa. The exact number of simulations for each set of taxa can be found in the Supplement Section 5.3.

Tree and sequence simulations are done using R packages ape [64] and phangorn [65]. We use the Jukes Cantor model [66] with a rate of 0.005 to generate alignments of 800 base pairs. We run BEAST2 with this model specified, a chain length of 2.5 million samples, discarding first 500, 000 as burn-in. Trees were sampled every 2, 000 iteration resulting in a tree set of 1, 001 trees. For larger analyses (taxa *≥* 100) we switched to 7 million samples (1 million burn-in) thinning to an output tree set of 2, 000 trees. A visual representation of the simulation workflow as well as the parameters used are displayed in Figure 1. Note that each simulation comprises one simulated alignment and one BEAST2 run on said alignment. To check convergence, we calculate ESS values using Tracer [67] and also use diagnostic tools from the RWTY package [68]. Both confirm convergence using the commonly used ESS threshold of 200 and following the plot interpretation provided by Warren *et al.* [68].

**Figure 1.**
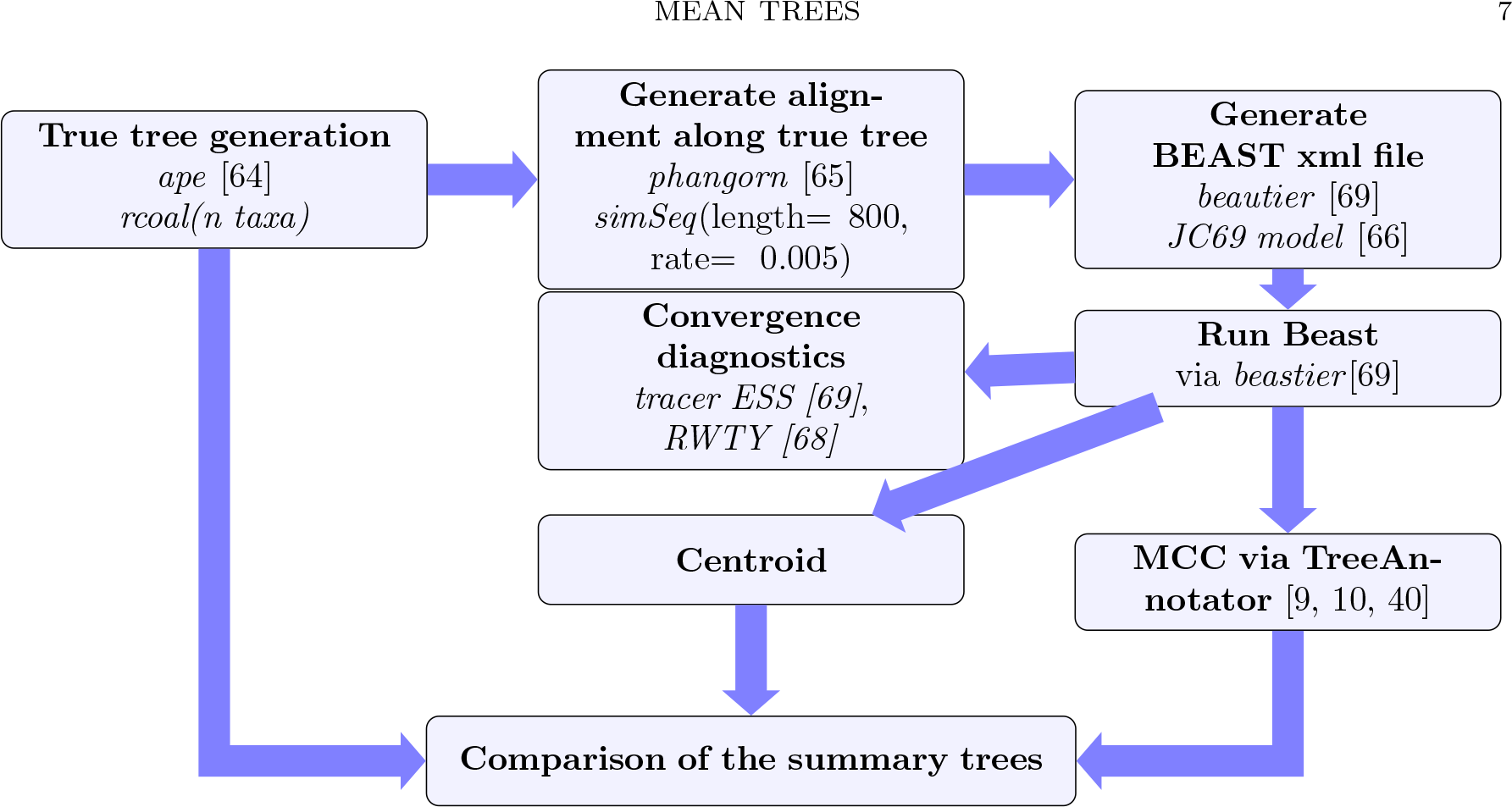
Flow chart of the simulation study

## 3. Results

In this section we first present results of our study comparing the approximate mean tree to the MCC summary tree, the current default summary method in a majority of applications, using an extensive range of error measures. We then analyse the behaviour of the log-likelihood function during the execution of our approximation algorithm. The “as expected” behaviour of the log-likelihood function in the RNNI treespace motivates our next investigation into the landscape of the log-likelihood function in different treespaces. We conclude the section by presenting applications of our method to three datasets in linguistics, phylogeography, and somatic evolution.

### 3.1. Mean trees in RNNI

In this section, we investigate how well our approach of approximating means of probability distributions over the RNNI treespace works in practice. Specifically, we establish various statistical and algorithmic properties of our method that are relevant for deploying our algorithm in practice. We start by comparing our approximate mean tree calculated by the centroid algorithm with the MCC tree, the current gold standard for summarising samples of posterior distributions from a BEAST or BEAST2 analysis. In this comparison we use the annotated centroid approximation (Section 2.2.2) and the MCC tree with its common ancestor annotation (Section 2.3).

Using simulations as described in Section 2.4, we compare the log-likelihood of our approximate mean tree with the log-likelihood of the MCC tree. The result is shown in Figure 2. The diversion off the diagonal (red) indicates that log-likelihood values are higher for the centroid approximation than for the MCC tree. The dots, each representing an individual simulation run, are coloured according to the RNNI distance between the two summary trees with darkness increasing with RNNI distance. Curiously, the darker dots are further away from the diagonal. We investigate this phenomenon in Section 3.2.

**Figure 2.**
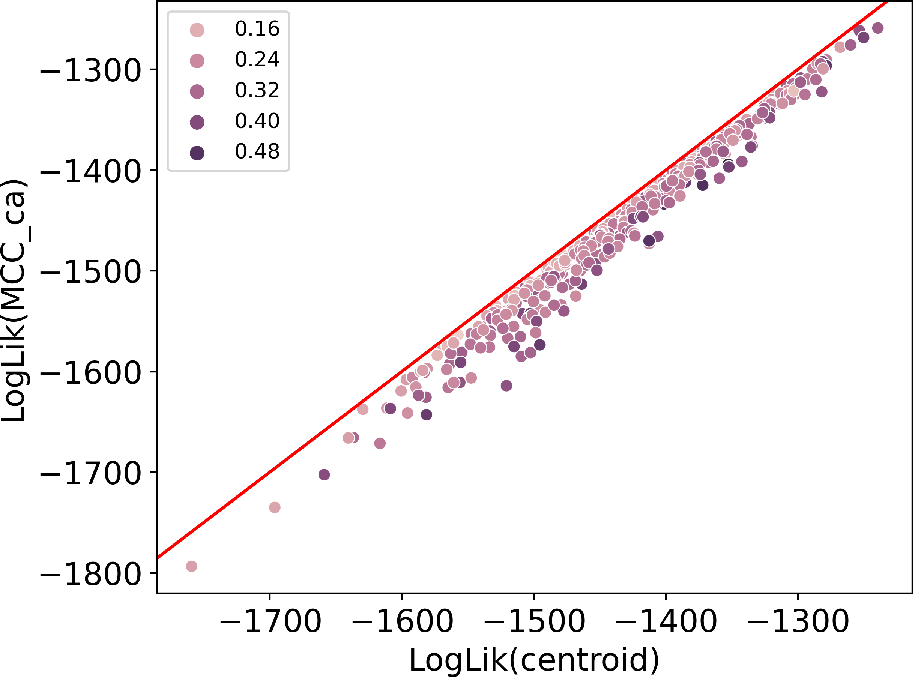
Comparing log-likelihood values of MCC trees with our centroid approximations. The data points are colour coded by their normalised RNNI distance (RNNI distance between the two summary trees divided by maximum possible RNNI distance). The red diagonal line shows the identity of the log-likelihood values.

A one-sided Mann-Whitney-U test [70] (as implemented in [71]) suggests that the distribution of log-likelihood values for centroid trees has a greater mean value than the one for MCC trees (*p*-value of 0.00013, indicating a significant difference in the means log-likelihood values). These results imply that our centroid approximation effectively provides a tree with higher likelihood than the MCC tree, making it a more accurate approximation of the mean tree and hence a better summary of a tree sample.

In addition to this likelihood comparison, we consider a number of other measures to compare the centroid approximation to the MCC tree. A summary of the results is presented in Figure 3, where all measures compare the two summary trees (centroid and MCC) to the true tree used for the simulations. More detailed figures can be found in Supplement Figure 16. The first two measures, clade age error and clade rank error [40], compare the height of clades in the respective summary tree to the true tree, and for both errors our mean tree approximation outperforms the MCC tree. The results for the clade rank error imply that the underlying ranked tree of the MCC tree is not as similar to the true tree as the ranked tree resulting from the Centroid algorithm. The result for the clade age error is surprising, as it suggests that our new method of annotating the ranked mean tree works well, even though it only takes heights of nodes into account regardless of the leaf labels below it. We also use the RNNI metric, where the approximate centroid tree is a more accurate approximation than the MCC tree. We finally consider weighted and unweighted Robinson-Foulds distance between the summary trees and the true tree [52, 72]. Under the regular Robinson-Foulds metric the MCC tree is (unsurprisingly) closer to the true tree than the centroid approximation is, implying that the MCC tree captures certain clade-based topological features of a set of trees better. However, the weighted Robinson-Foulds distance to the true tree is always smaller for the centroid approximation, confirming that the centroid approximation in combination with our new branch length annotation method outperforms the MCC summary.

**Figure 3.**
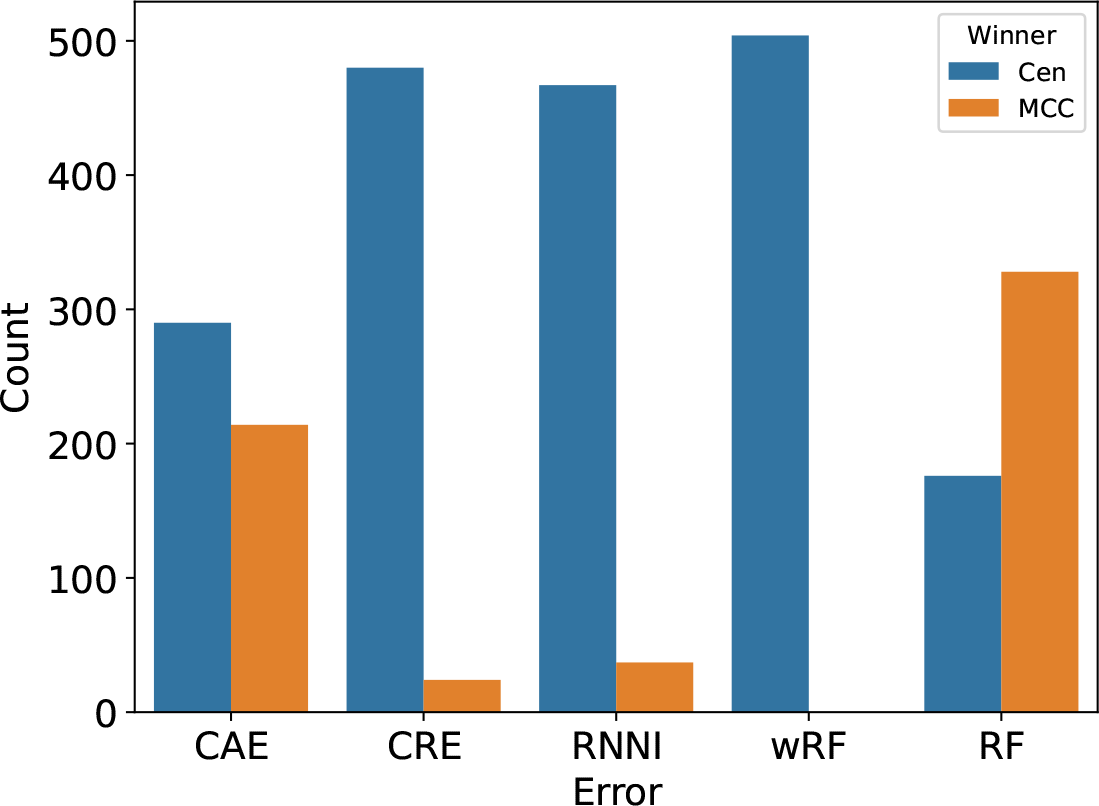
Comparison of the centroid approximation and MCC tree using different evaluation measures. The y-axis shows in how many simulations either MCC or the centroid approximation was superior. The error measures are clade ages error (CAE), clades rank error (CRE), RNNI metric, weighted Robinson-Foulds (wRF) and regular Robinson-Foulds metric (RF). All measures compare the summary trees with the true tree for each simulation.

We additionally analysed smaller datasets with 8–50 taxa, which show similar results (see Supplement Figure 17). Only for trees with fewer than 20 taxa, the centroid approximation and the MCC tree were mostly identical, which indicates that when the tree sample is well converged both summary methods are able to approximate the mean tree well. However, the difference of the two summary trees increases significantly when the number of taxa increases.

To disentangle the effect of our branch length annotation method (Section 2.2.2) from our estimation of the ranked tree topology, we investigated the impact of changing the branch length annotation method. Specifically, we compared log-likelihood values of the MCC tree topology annotated with the three default methods provided in TreeAnnotator with our newly presented method. The comparison demonstrates that there is no significant difference of log-likelihood values in the MCC trees returned by either of these branch length annotations. These results are visualised in the Supplement (Figure 15, Figure 14 and Figure 18). A detailed comparison for each of the error measures from Figure 3 can also be found in the Supplement (Figure 16).

Our method is implemented to run in practical time and is unrestricted in its exploration of the RNNI treespace to find a good candidate tree (see the Supplement Section 5.6 for details), improving over existing geometric mean approaches that suffer from long runtime and only explore the respective treespace partially, often restricted to a small set of input trees [39, 40].

### 3.2. Likelihood and posterior distributions over the RNNI treespace

In this section we further investigate the relationship between the geometry of the RNNI treespace and the likelihood function. Specifically, we are looking into the changes of the likelihood function during execution of the Centroid algorithm. Recall that at every iteration of the algorithm the SoS value from the running tree to the tree sample decreases. The result is depicted in Figure 4, where we plot the log-likelihood value against the SoS distance over the iterations of the Centroid algorithm (Algorithm 1). We can see that while the algorithm minimises the SoS value of the approximate mean tree, the log-likelihood value increases. Hence, the Centroid algorithm finds a tree with higher and higher log-likelihood while only using the geometry of the RNNI treespace. This connection between the geometry of the space and the behaviour of likelihood functions indicates the appropriateness of this treespace for statistical analyses of phylogenetic time trees.

**Figure 4.**
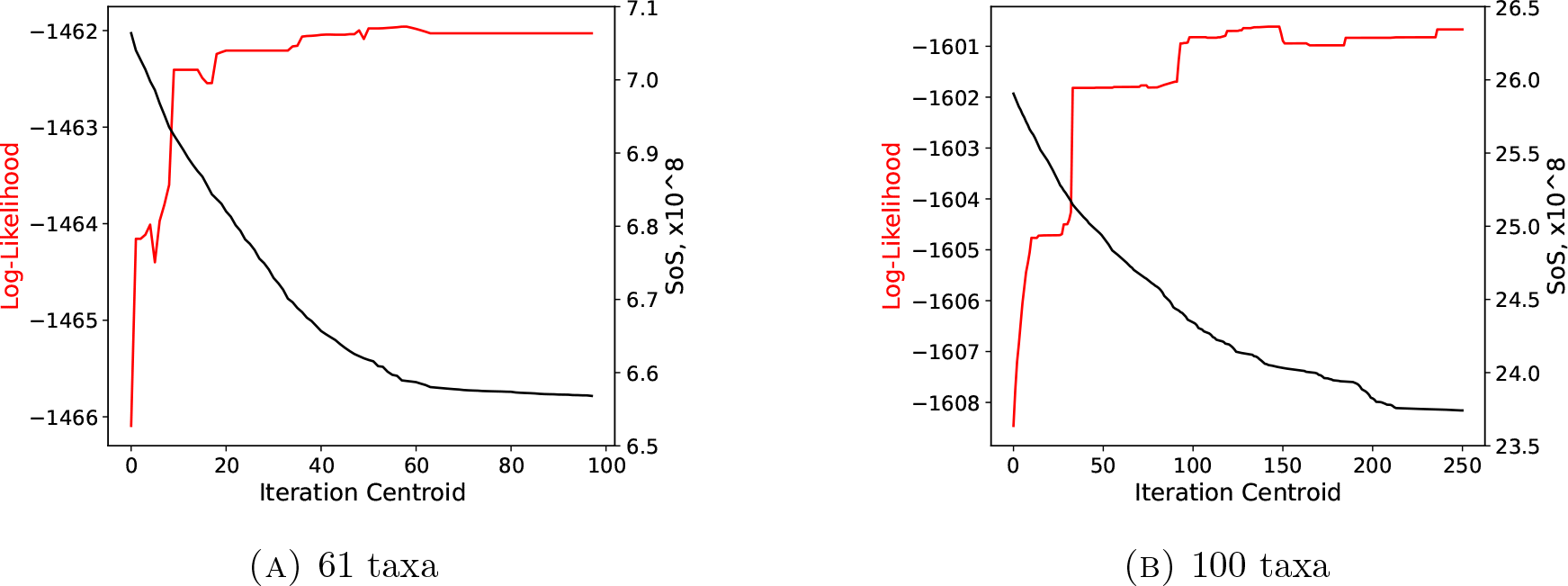
Log-likelihood and SoS values for two different Centroid runs on simulated data sets with varying number of taxa.

To further investigate the relationship between the geometry of the RNNI treespace and the likelihood function, we investigate the correlation between the SoS and log-likelihood value of a tree. We found a negative correlation with both the Pearson and the Spearman tests (as implemented in [71]) ranging between *−*0.4 and *−*0.6 among our simulated data sets, see Supplement for a visualisation of these values. In Figure 5 we compare the SoS and log-likelihood values for every tree within one simulated data set on 50 taxa. According to this negative correlation we have found another indication that the RNNI space is consistent with statistical intuition, i.e. the tree at the peak of the log-likelihood distribution should also be a minimum for the SoS values.

**Figure 5.**
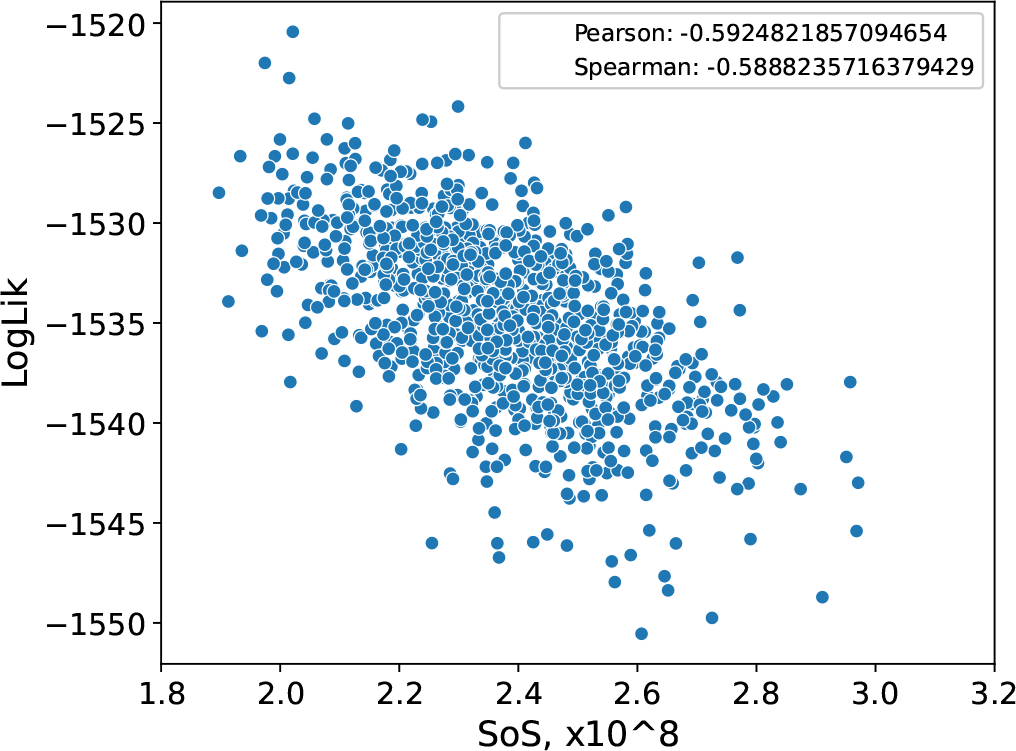
Log-likelihood values against SoS values for a set of trees from a simulation on 50 taxa. The top right legend contains the Spearman and Pearson correlation coefficient. Both indicating a negative correlation with a value of roughly *−*0.6 in this case. See Supplement for correlation coefficients among all simulated datasets.

Finally, we investigate how continuous the log-likelihood function is in different treespaces. The intuition here is that local changes in a treespace should correspond to local changes in likelihood and posterior probabilities of the corresponding trees. A degrees of continuity (or “smoothness”) is a desirable property for the commonly deployed hill climbing tree search algorithms in both Bayesian and maximum likelihood frameworks.

Using a simulated dataset as described in Section 2, we compare the (relative to the diameter) RNNI distance between pairs of trees to the difference between their log-likelihood and posterior probabilities. We also repeat this comparison for the BHV, Kendall-Colijn [22], and Robinson-Foulds metrics. We note that these metrics are designed for standard phylogenetic (rather than time) trees, and it has been shown [21] that adapting standard metrics to time trees is non-trivial. We found that both log-likelihood and posterior probabilities are “smooth” in the RNNI and Robinson-Foulds spaces but not in BHV and Kendall-Colijn. The result for BHV and the RNNI treespace is shown in Figure 6, and results for Robinson-Foulds and Kendall-Colijn metrics can be found in Supplement Fig-ure 20.

**Figure 6.**
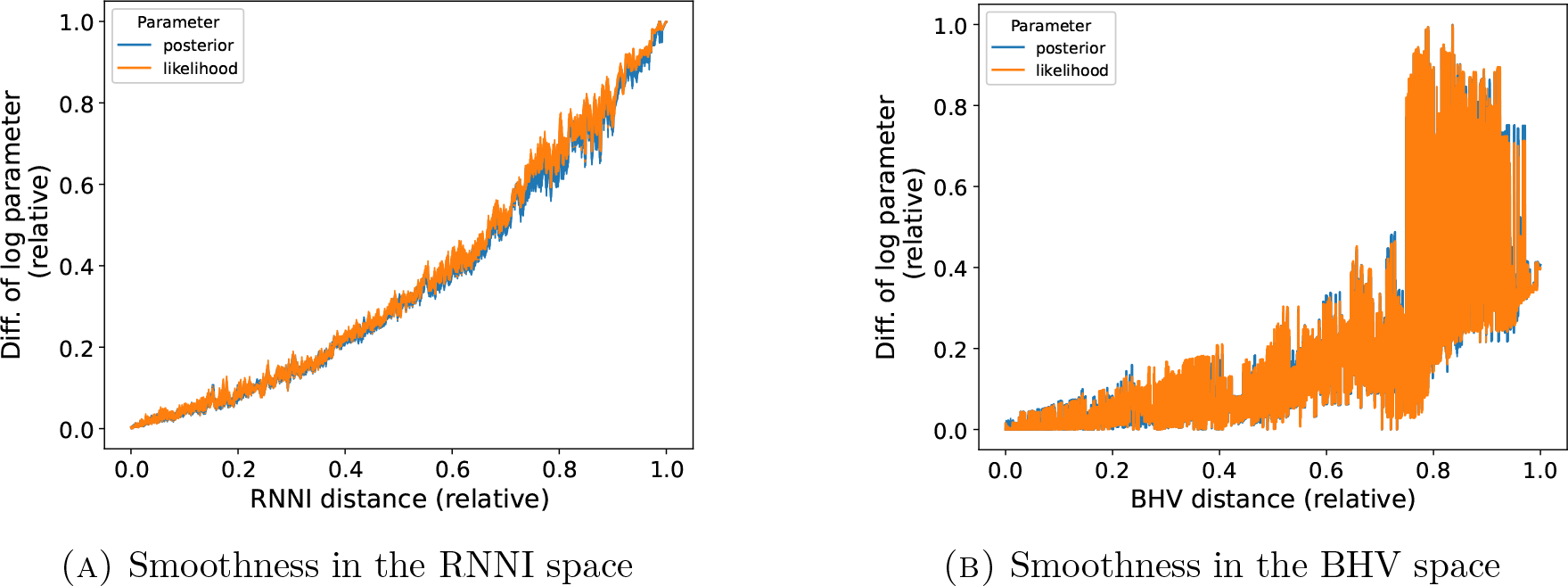
Assessing the smoothness of probability functions over the RNNI and the BHV treespace.

### 3.3. Real Data examples

In this section, we apply our method to three problems in applied phylogenetics. These include natural language evolution [73], molecular evolution [74], and cancer evolution [75]. In all three cases our method provided additional insights into (if not alternative hypotheses about) the corresponding evolutionary processes, demonstrating the value of our approach in practice.

#### 3.3.1. Dravidian languages

Our first application is in evolutionary linguistics. Specifically, we analyse the Dravidian language data set published by Kolipakam *et al.* [73]. The Dravidian languages are a diverse family of languages spoken by about 250 million people predominantly across southern and central India. The original study aims to provide a time-depth estimation of divergence between languages and a reconstructed evolutionary history of the language family. These results contribute to a deeper understanding of the historical development and diversification of the Dravidian language family, a topic of ongoing research with many unanswered questions.

For our analyses we took a BEAST2 sample of trees generated in the original study [73], applied our method, and compared the result to the published (MCC) tree. As it can be observed in Figure 7 the topology of our (centroid) tree is slightly different to the MCC tree (highlighted in yellow in the figure). In particular, our tree splits the sister relationship of the two languages Telugu and Koya placing Koya as a descendant of Telugu. As this clade of the South II languages is of specific interest in [73], where further investigation into this part of the tree is suggested, our result can serve as an argument in favour of this particular placement of the two languages.

**Figure 7.**
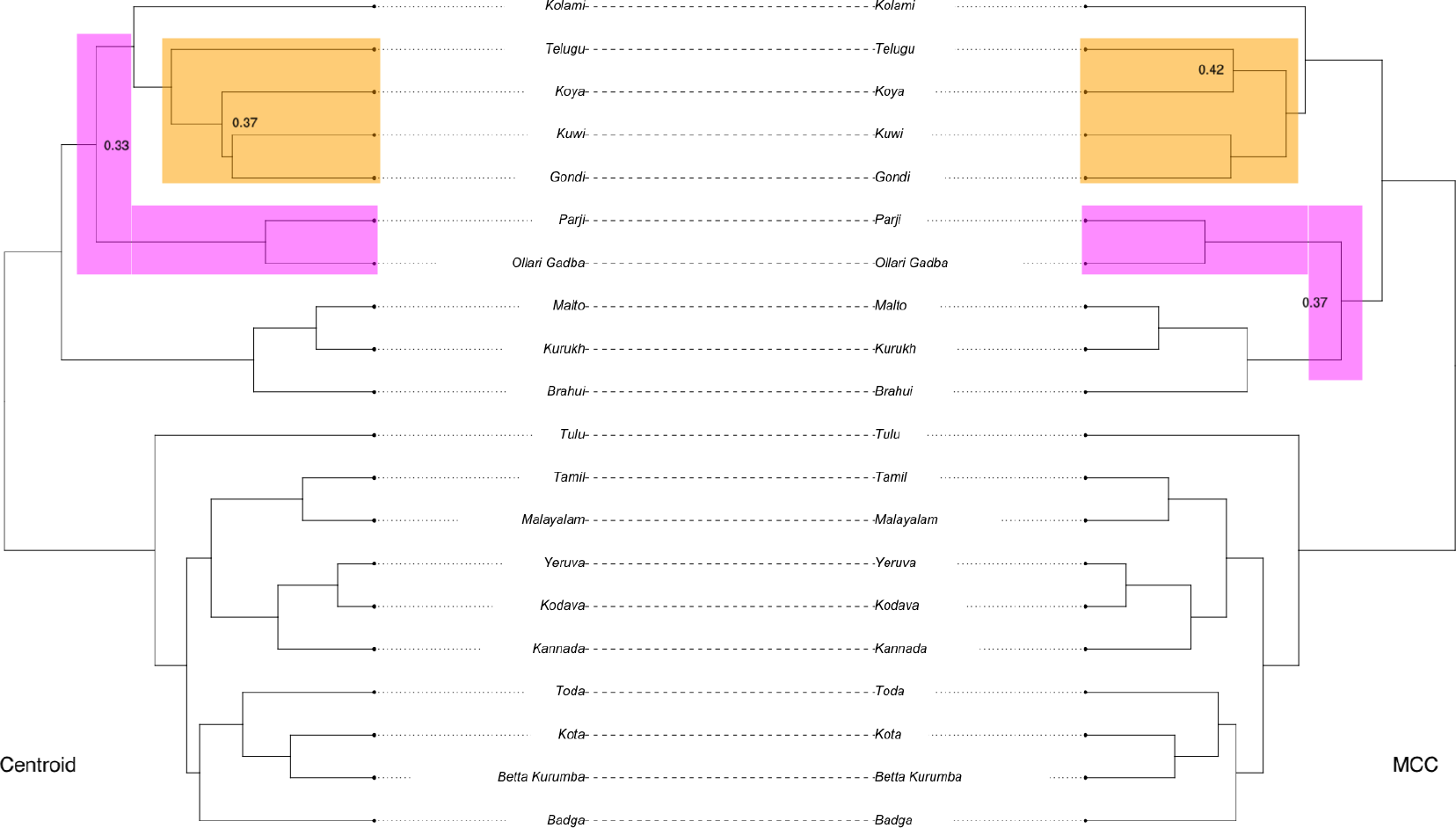
Comparing the MCCtree (right) to the centroid approximation (left). Highlighted clades have a changed topology or position in the centroid tree versus the MCCtree

The other difference between the two trees (highlighted in pink in Figure 7) is the degree of separation between the three Central languages (Parji, Ollari Gadba, and Kolami). Specifically, the two languages Parji and Ollari Gadba are closer to the Kolami language in the centroid tree whereas these form a distinct clade with the three North languages (Kurukh, Malto, and Brahui) in the MCC tree. This difference is consistent with the proximity network provided in Figure 2 in [73].

These discrepancies may be caused by convergence issues or a multimodal posterior distribution. Thus, in the context of data and discussion presented in [73] we conclude that the centroid topology provides valuable insights into the data and evolutionary process under consideration.

#### 3.3.2. Weevils

Our second application falls within the study of molecular evolution, specifically investigating in situ speciation on the islands Mauritius and Reunion of Cratopine weevils. The study applies phylogenetic and biogeographic methods to analyse genetic data from multiple weevil species and reconstruct their evolutionary relationships and dispersal patterns. The results of the study provide new insights into the processes of community assembly and diversification in a species-rich island radiation, and highlight the role of historical and ecological factors in shaping the evolution of these weevils. Here we are interested in the specific differences of topologies between the MCC and our centroid tree and their implications for the data set.

As noted by Kitson *et al.* [74] the position of the *C. nigrogranatus* in the phylogeny is unresolved. We find that its position is indeed different for the two trees, see Supplement Figure 21. Moreover, we find that the position of one of the Rodrigues clades containing *C. virescens* and *C. viridipunctatus* has changes its position within the larger subtree. It does in fact not move any closer to the second clade thought to colonise Rodrigues keeping the question of multiple colonizations of Rodrigues open. This clade is now located in the middle of the clade 2b (see Figure 2 in [74]), suggesting a different lineage for these species and the respective colonization process of the islands.

#### 3.3.3. Cancer evolution

Our third application is within the domain of cancer evolution, specifically the evolution and spread of colorectal cancer cells within a patient. Alves *et al.*[75] presented tumour clones computationally inferred from whole-exome sequencing data and analysed the history of cancer cells evolving and spreading to different organs over time. The results from this study provide important insights into evolutionary dynamics of cancer within a single patient by identifying tumour demographics and colonization patterns in a defined time-frame. Here we repeat the original analysis using our centroid method and compare the result with the published (MCC) tree.

The main difference between the two trees is a change of the subtree topology corresponding to the liver, colonic lymph node, and hepatic lymph node metastases clade (taxa D, F, J, I, R, L, G in Figure 8). In the MCC tree the corresponding subtree is a fully balanced topology, in which the liver cells are grouped with colonic and hepatic lymph node cells as cherries in the tree. In the centroid tree on the other side, this clade is resolved as a caterpillar topology where all the liver cells are grouped as single leafs one after another. Because the tree shape can be informative of the type of evolutionary process [76, 77] and e.g. can be used to distinguish among different epidemiological scenarios, the difference between the MCC and centroid tree shapes can point at different evolutionary processes driving cancer development in this clade. In Figure 8 we also highlight other less significant differences in branch lengths and timing of nodes.

**Figure 8.**
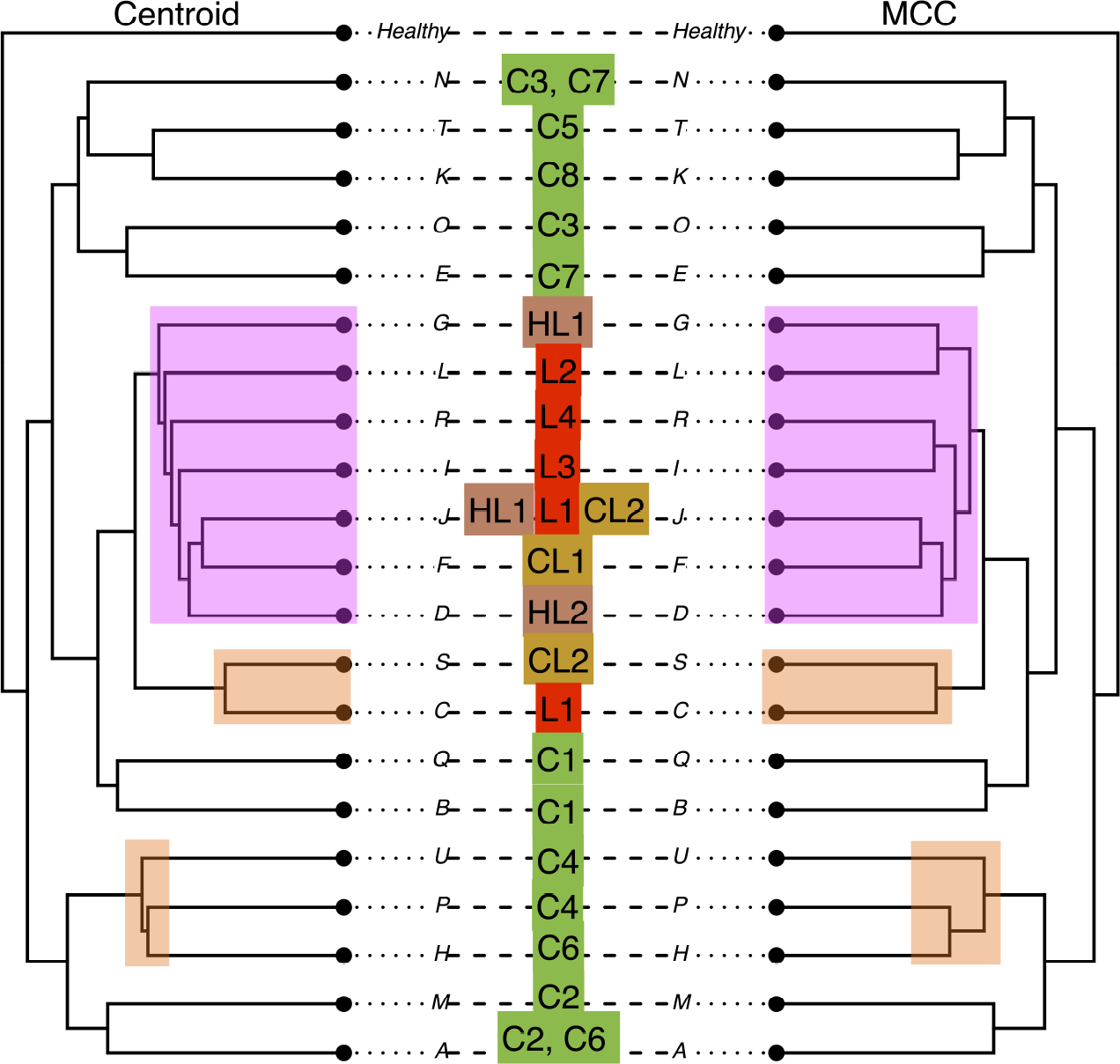
Highlighted differences in the two trees for the analysis of cancer data.

## 4. Discussion

The RNNI treespace has been developed specifically for phylogenetic time trees and it is the only treespace, to the best of our knowledge, that is based on local tree rearrangement operations with efficiently computable distances and paths. In this paper, we study the question of whether this property enables the RNNI space to be suitable for developing probability theory and statistics over the space of phylogenetic time trees. We show a number of promising results that collectively suggest that the answer to this question is “yes, but a significant amount of work is yet to be done”. For example, we are able to demonstrate a strong connection between RNNI as a metric space and log-likelihood as a probability distribution over this space. This connection came in the form of a smoothness property of the log-likelihood functions over the RNNI space, which we believe is a desirable property of a treespace. Further, we developed an algorithm, Centroid, for approximating means of probability distributions in this treespace. As a practical consequence of this algorithm, we demonstrated its excellent performance at summarising samples from posterior distributions in the full Bayesian framework, consistently outperforming the state of the art approach, the MCC method. This result has concrete consequences for data analysis, and we demonstrated this by applying our algorithm in three areas including natural language, molecular, and cancer evolution. In all three cases, the Centroid tree was different from the MCC tree reported in the original study. Although in all cases our summary tree was compatible with the discussions in the respective papers, it supported some of the alternative hypotheses the authors considered originally. So we conclude that our method has immediate practical utility.

Having scratched the surface of what we believe can be achieved with a local rearrangement based treespace with efficiently computable distances and paths, we finish this paper by outlining some of the possible ways forward. First, we summarise the properties of the RNNI treespace that we believe are fundamental for a treespace to possess in order to be suitable for statistical analyses. We intentionally leave the statements of these properties mathematically informal as their formalisation would inevitably leave some potentially fruitful approaches to developing statists over treespaces out, which is something we intentionally avoid. For example, we would like to avoid requiring the treespace to be a metric space.

1. The treespace is local, that is, the notion of a (local *δ*-) neighbourhood is well-defined and all trees within a neighbourhood are “similar”; computationally, neighbourhoods need to be efficiently computable/samplable. For RNNI this is enabled by tree rearrangement operations.
2. The notion of “betweenness” is well-defined, that is, it is possible to find a weighted average between two (or more) trees, and the average tree is “similar” to the trees that are being averaged; computationally, these need to be efficiently computable or approximated. For RNNI this is enabled by shortest paths satisfying the cluster property and the property that all trees on shortest paths are fully resolved.
3. Probability distributions (of importance) over the treespace are “smooth”, that is, similar trees have similar probabilities; computationally, the likelihood functions need to be efficiently samplable. For RNNI this is enabled by standard phylogenetic likelihood functions being efficiently computable and local with respect to tree rearrangement operations.

This list is certainly not exhaustive and our main reason for providing it here is to encourage a discussion of desirable properties a treespace should possess depending on the use case. We are hopeful that this discussion would enable a more purposeful treespace design and would guide phylogenetic practitioners whenever a choice needs to be made regarding what treespace to use.

Second, we provide a list of mathematical problems that we believe would be important to solve.

1. Given a set *S* of *m* trees in RNNI such that *m > n* (the number of leaves):

a. How many mean trees can *S* have?
b. Can a mean tree of *S* be computed in polynomial in *m* time?
c. Can all mean trees of *S* be computed in polynomial in *m* time, assuming the number of mean trees of *S* is polynomial in *m*?

Related to this is the question of whether the running time of the Centroid algorithm can be improved, that is, whether there exists a faster algorithm for approximating mean trees in the RNNI space. Although the running time of the Centroid algorithm for big trees is still significantly higher than that of the MCC algorithm, there is nothing to suggest that our implementation is optimal.

We, in fact, have carried out several computational experiments aimed at understanding the number of mean trees, as well as scenarios when the Centroid algorithm fails to find a mean tree and the impact of the starting tree on the algorithm’s performance. These are presented in Supplement Section 5.1.

(2) Given the mean tree in the RNNI space:

a. Is the branch length annotation we use for ranked trees statistically consistent?
b. Is there a better way to annotate branch lengths?

Although potentially statistically inconsistent, our annotation method works well in practice. Either way, we have little doubt that our method can and should be improved.

## Acknowledgement

This work was partially supported by Royal Society Te Aparangi through a Rutherford Discovery Fellowship (UOC1702) and a Marsden grant (21-UOC-057), and by Ministry of Business, Innovation, and Employment of New Zealand through an Endeavour Smart Ideas grant (UOOX1912) and a Data Science Programmes grant (UOAX1932). We are grateful to David Posada and Joao Alves for providing us with additional datasets and helpful comments. We thank Alexei Drummond for advice and helpful suggestions about the presentation of the results. We also thank Mareike Fischer, Russell Gray, Remco Bouckaert, Allen Rodrigo, and the members of the bioDS lab for their feedback on earlier versions of this paper.

## 5. Supplement

### 5.1. Centroid properties

#### 5.1.1. Finding global optima

To conduct a test of whether or not our algorithm is able to recover global optima as presented in Figure 9, we implemented the following approach:

- Generate all trees of the treespace (up to 7 taxa)
- Generate a random tree set in this space
- Compute the SoS for all trees and save the global optimum of SoS
- Start the Centroid^1^ algorithm from every tree in the tree set
- Combine all solutions and choose the best one (minimal SoS)
- Check if it is a global optimum

**Figure 9.**
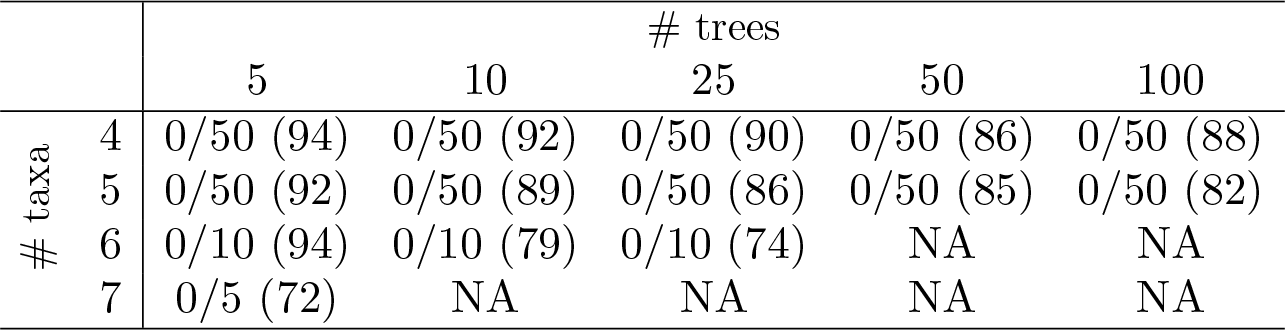
Displays the results of tests to see if the Centroid algorithm finds a globally optimal solution. The row index shows the number of taxa and the column index the size of the randomly generated tree set. An entry *x/y* (*z*) displays the number of times *x* the algorithm did not find a globally optimal solution out of *y* different randomly generated tree sets. The value (*z*) displays the percentage of trees in the tree sets that when used as a starting tree found a global optimum. *NA* are not conducted tests because of the increasing time complexity.

We conclude that the algorithm is likely to always find a global optimal solution when the starting tree is chosen correctly. However, starting the algorithm from every tree in the set and also following all paths that decrease the objective function, i.e. the SoS value, already takes a long time in these small treespaces and tests on 6 and 7 taxa were only conducted on very small tree sets (see NA in Figure 9).

#### 5.1.2. How many global optima are there?

We further investigate these globally optimal solutions, specifically in the context of how many such optima exist and also whether or not these optima form a connected subgraph in the treespace. The findings presented in Figure 10 lead us to formulate the following conjecture

**Figure 10.**
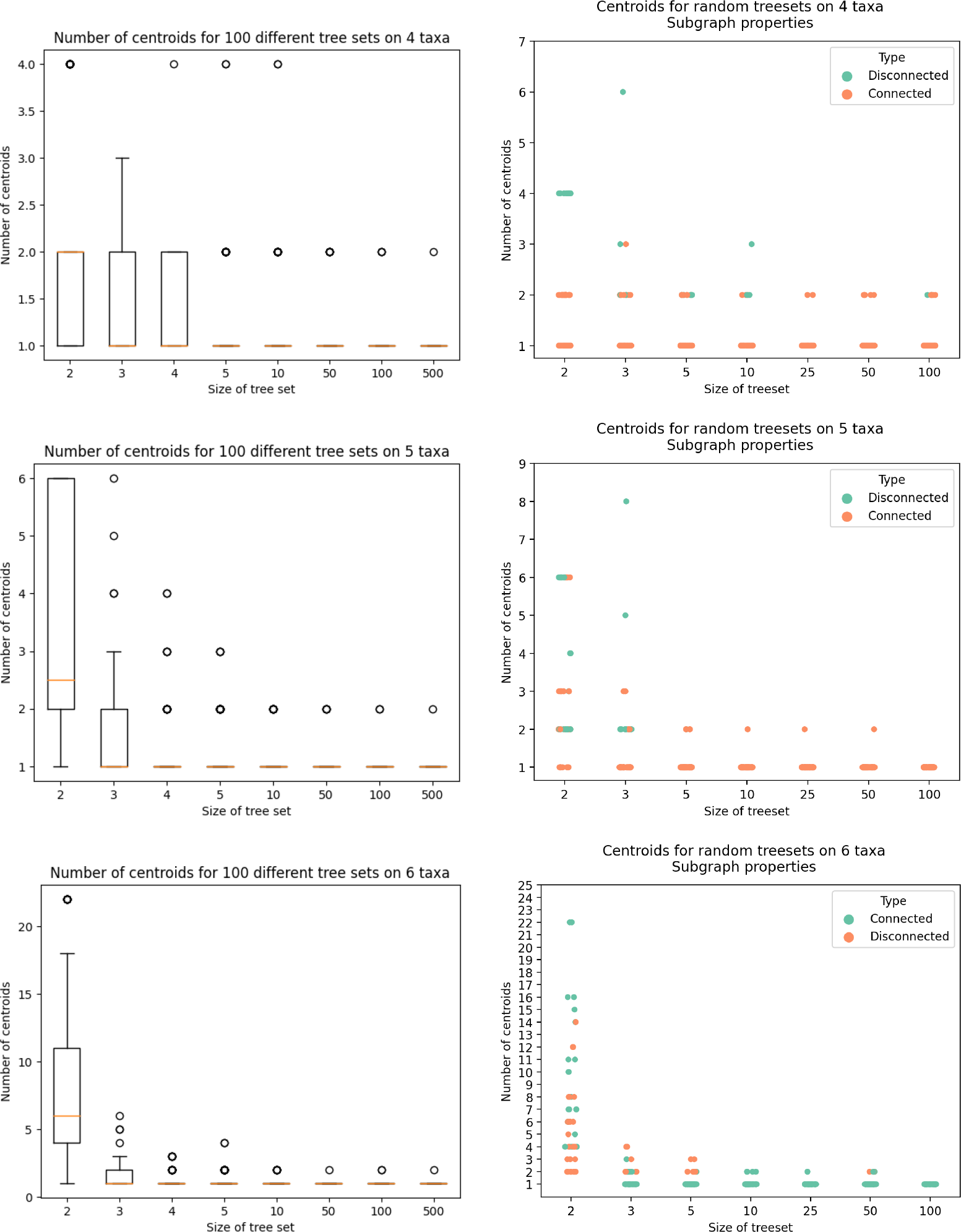
Evaluating number of centroids and their connectedness on small taxa RNNI treespaces. Left column plots are for 100 randomly generated tree sets of different sizes and the right column plots are from 50 randomly generated tree sets.

**Figure 11.**
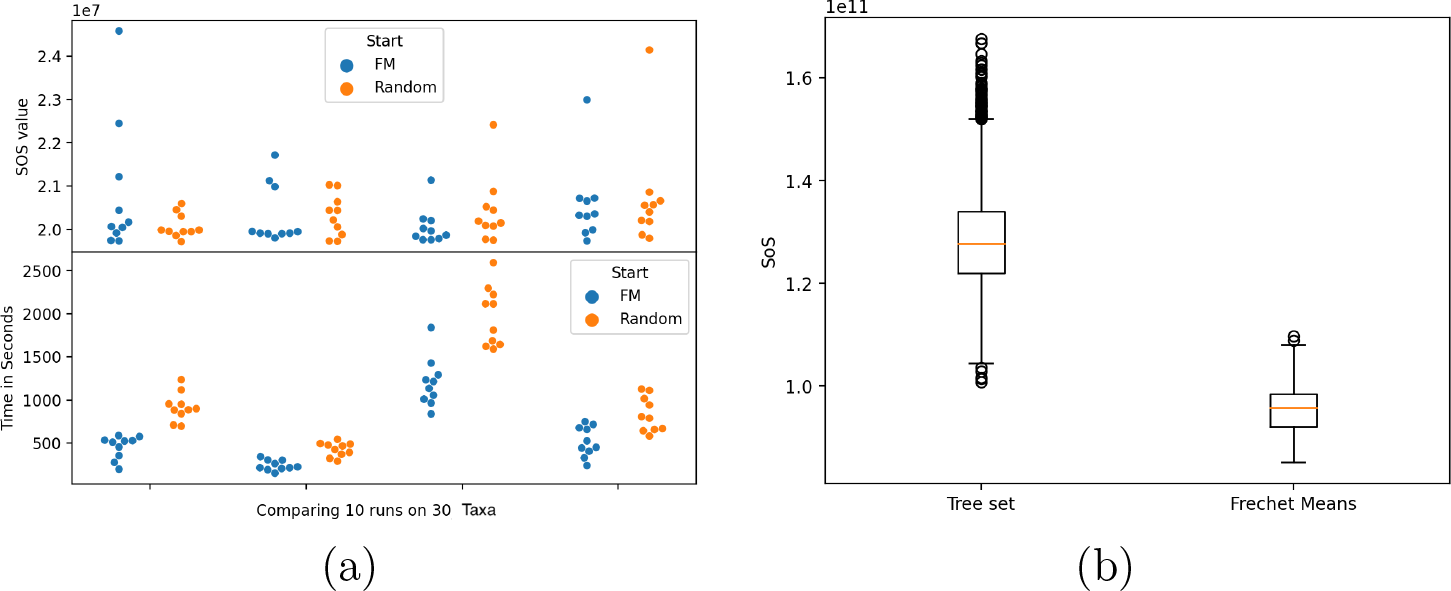
(a) Comparing 4x10 different executions of the Centroid algorithm with a random starting tree from the set vs. a tree computed by the Starting Tree algorithm (FM in the plot) on a dataset with 30 taxa. The top half displays the SoS value of the returned tree of the algorithm and the bottom shows the runtime of a Centroidcomputation using either of the starting trees. (b) Comparing the SoS value of all samples in a set of trees versus the SoS value of starting trees computed by the Starting Tree algorithm on a dataset with 188 taxa. Due to the random choice of trees in the Starting Tree algorithm there is some variance in the returned tree.

**Conjecture 5.1.** For a sufficiently large tree set there exists a unique centroid tree.

Starting Tree *algorithm.* Pseudocode for the variation of Strums algorithm [46] adapted to the RNNI treespace. For geodesics we use the shortest path within the graph that is computed by the findpath algorithm [26, 62].

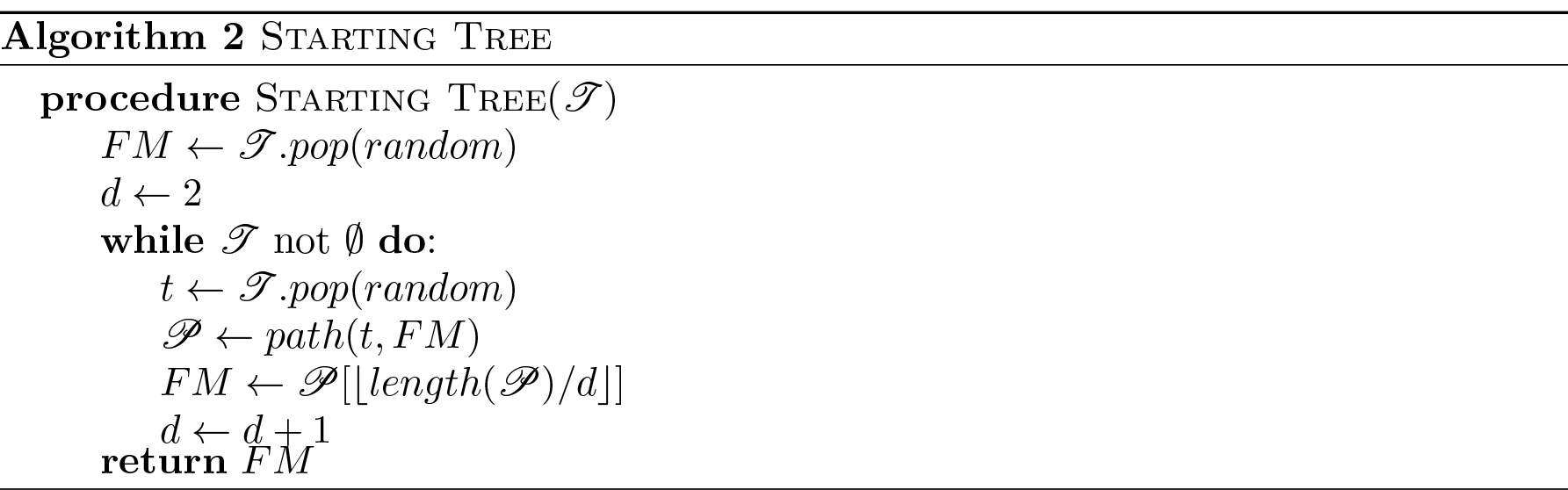

In addition we run some tests on smaller dataset to compare the output of this algorithm with any tree that could be picked from the original tree set. It turned out that a starting tree computed by this algorithm is consistently below the expected SoS value one would get by picking a tree in the given set of trees. This result is consistent with geometrical intuition in a treespace.

### 5.2. Error Measures

Most of these definitions for error measures can be found in [40].

*Clade ages error -* CAE. The *clade ages error* CAE, the sum of differences between clade ages, for a tree *T* and a reference tree *T_R_*is defined as

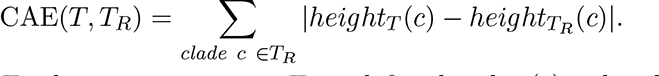

For a clade *c* in *T_R_* that is not in a tree *T* we define *height_T_* (*c*) = *height_T_* (*mrca*(*c*)). It is important to note that this error is not symmetrical.

*Clades missed error -* CME. The *clades missed error* CME counts the number of clades present in the reference tree *T_R_* which are not in *T* :

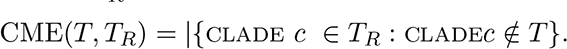

This error is exactly half of the RF distance [40].

*Clades called error -* CCE. The *clades called error* CCE scores +1 for correct clades and *−*1 for incorrect clades in a tree *T*, compared to a reference tree *T_R_*:

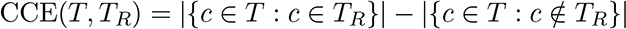

*Clade rank error -* CRE. The *clade rank error* CRE is a deviation of the clade ages error CAE. It is the sum of differences between clade ranks, therefore this error measure is specific for ranked phylogenetic trees:

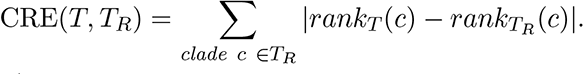

Analogue to the CAE, the rank of a clade *c* that is not in a tree is defined as the rank of the most recent common ancestor of that clade. This error is also not symmetrical.

*Tree metrics.* The considered tree metrics and their used abbreviations.

- Robinson-Foulds – RF
- weighted Robinson-Foulds – wRF
- Path difference – PD
- weighted path difference – wPD
- Branch Score difference – KF
- RNNI distance – RNNI

*Log Likelihood value.* For the log-likelihood value we use the *pml* function from the R package *Phangorn* [65].

### 5.3. Simulation Study - number of simulations

The following table Table 1 shows the actual number of simulations that were conducted. The reason for the ”weird” uneven numbers are solely for the ease of implementation and due to increasing complexity and time consumption for higher numbers of taxa.

**Table 1.** Overview of the data used in the following comparison

|  |  |  |  |  |  |  |  |  |  |  |  |  |  |  |  |  |  |
| --- | --- | --- | --- | --- | --- | --- | --- | --- | --- | --- | --- | --- | --- | --- | --- | --- | --- |
| n taxa | 50 | 51 | 55 | 60 | 65 | 70 | 75 | 80 | 90 | 100 | 120 | 125 | 140 | 150 | 175 | 200 | Total |
| k simulations | 50 | 10 | 50 | 50 | 50 | 50 | 50 | 51 | 50 | 20 | 8 | 13 | 15 | 15 | 12 | 10 | 504 |

### 5.4. Simulation Study - Convergence Checks

We checked for convergence of our simulated data sets with the current state of the art tools. For this we checked that the ESS values are all well above 200 with the Tracer tool. Additionally we used the R package RWTY to check different diagnostics.

### 5.5. Conversion from branch lengths to ranking

Because the trees returned by a BEAST analysis are time trees the real times have to be converted to the discrete ranks of the RNNI trees. There are different ways of calculating the corresponding ranked tree. For these results we used a top down approach for conversion where the ranks are inferred by the corresponding t-space coordinates [21], similar to how we annotate a tree with branch lengths Section 2.2.2.

### 5.6. Running time of Centroid algorithm

Because of the problems with long running times of [39, 40] previous attempts at implementing a geometric mean based method we evaluate our implemented approximation. We found that due to our use of the greedy path heuristic in combination with the efficiently computable RNNI distance that the algorithm is running in reasonable time. Unlike the algorithm presented in [40] (named minimum distance tree) our method is also not restricted on a subset of trees and able to explore all of treespace. These tests are performed using 8 threads across two eight-core (16 SMT threads) Intel(R) Xeon(R) Gold 6244 CPUs in a NUMA system and a summary is provided in Figure 12.

**Figure 12.**
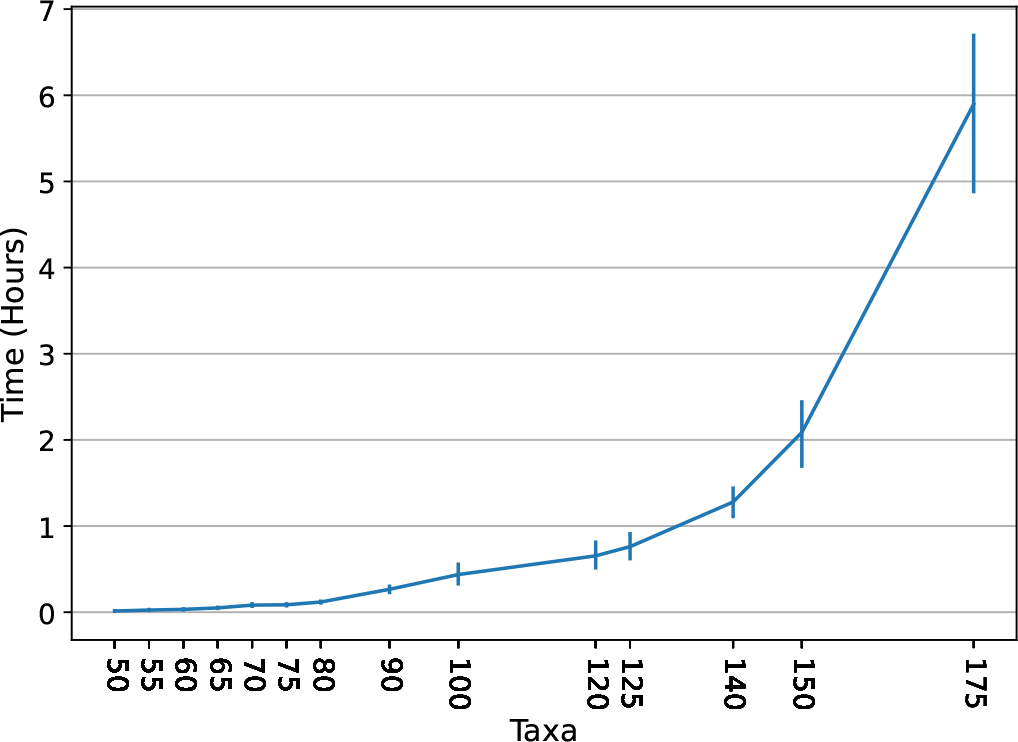
Evaluating the running time of the Centroid algorithm as implemented in [78].

### 5.7. Correlation of SoS and log-likelihood

As stated in the main paper we found negative correlation between the log-likelihood and the SoS value of trees. The following plot (Figure 13) visualizes the correlation coefficients among all datasets separated by the number of taxa.

**Figure 13.**
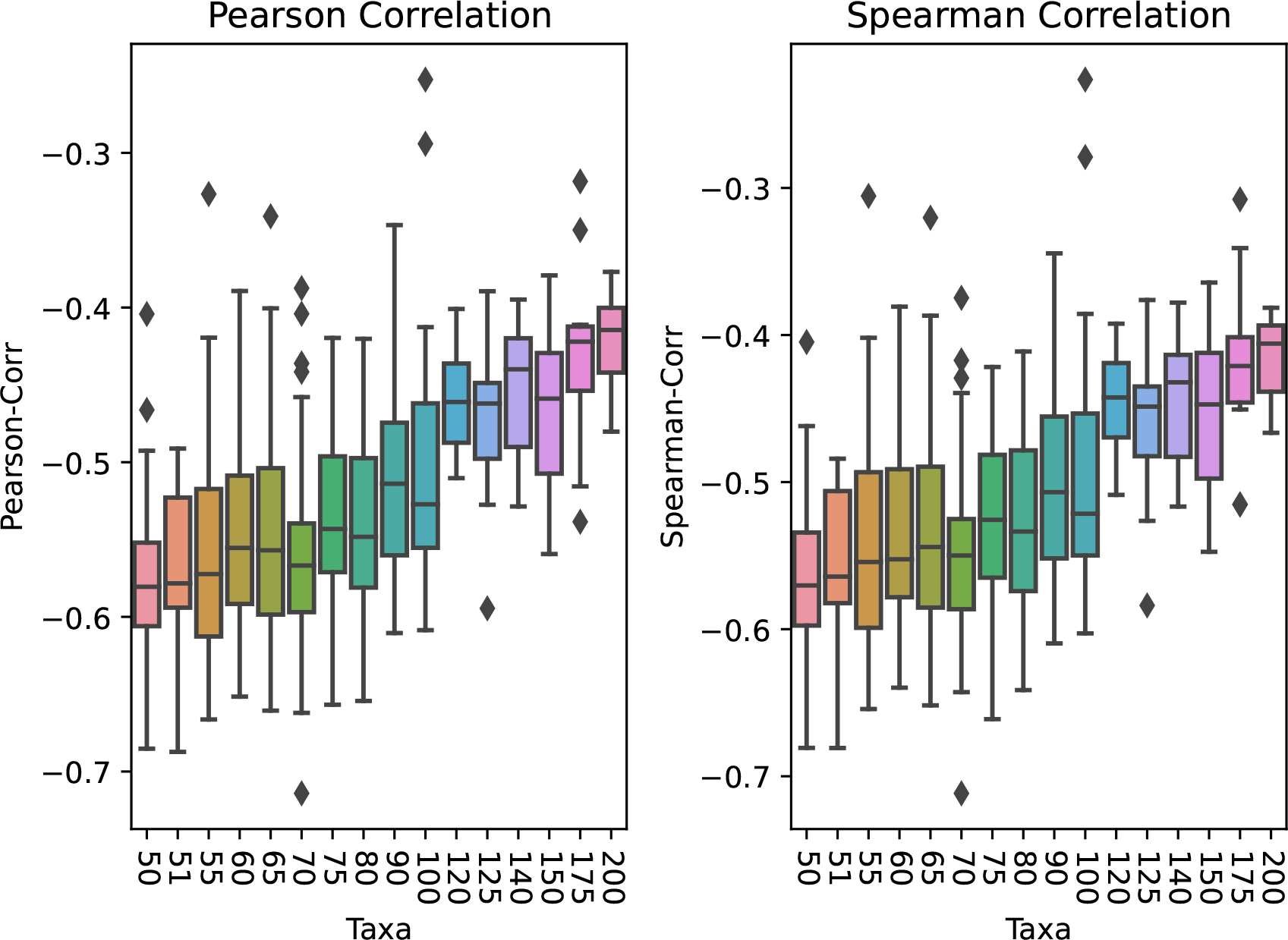
Pearson and Spearman correlation coefficients for all simulated datasets, separated by number of taxa.

### 5.8. Different MCC branch length annotations

The *TreeAnnotator* program provides three different branch length annotation options. The first option uses the ages as they are in the tree picked from the posterior sample. The Second and third option com-pute mean and median clade ages, respectively, for each clade based on their age in all trees from the posterior sample. Both of these annotations can result in negative branch lengths, because the average height of one clade in the MCC tree may be higher than the average height of its parent [63]. The fourth option computes the average most recent common ancestor time for each clade, therefore avoiding negative branches being annotated [40, 63]. We compare the three different tree annotations for the MCC tree (we do not consider the method of keeping the branch lengths from the picked tree here because we think it is vital to use all the information of an analysis and not just pick one specific tree) to choose the best one for the comparison with our new method. The three differently annotated MCC trees are compared based on their log-likelihood values, as displayed in Figure 15. The plot shows slightly worse log-likelihood values for the common ancestor option than for median and mean branch length annotation. We conduct a Mann-Whitney-U test [70] on the log-likelihood values and found no significant difference in the log-likelihood values of these three annotations. To investigate whether our method, presented in Section 2.2.2, of annotating branch lengths improves the MCC summary tree, we took the ranked tree topology produced by the MCC, annotated it using our method, and compared the result with the mean MCC branch length annotation as implemented in TreeAnnotator. We found no significant difference between the log-likelihood of these two trees (visualized in Figure 14) and therefore, we only compare the centroid approximation to a MCC tree using either of the three already available branch length annotations. However, as both mean and median branch length annotation result in negative branch lengths in some of our simulations, we compare our centroid approximation with the common ancestor annotation of the MCC tree.

**Figure 14.**
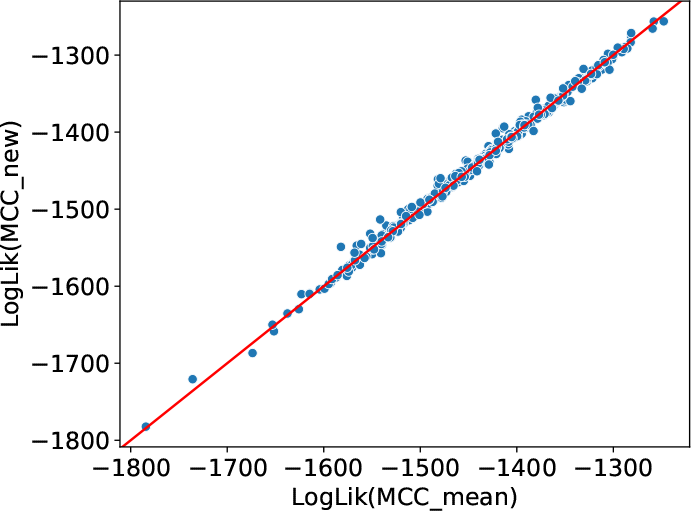
Comparing the log-likelihood values of the mean MCC branch length annotation versus our presented way of annotating a ranked tree (Section 2.2.2). This plot displays slightly more deviation of the red diagonal line than present in any of the three comparisons of Figure 15.

**Figure 15.**
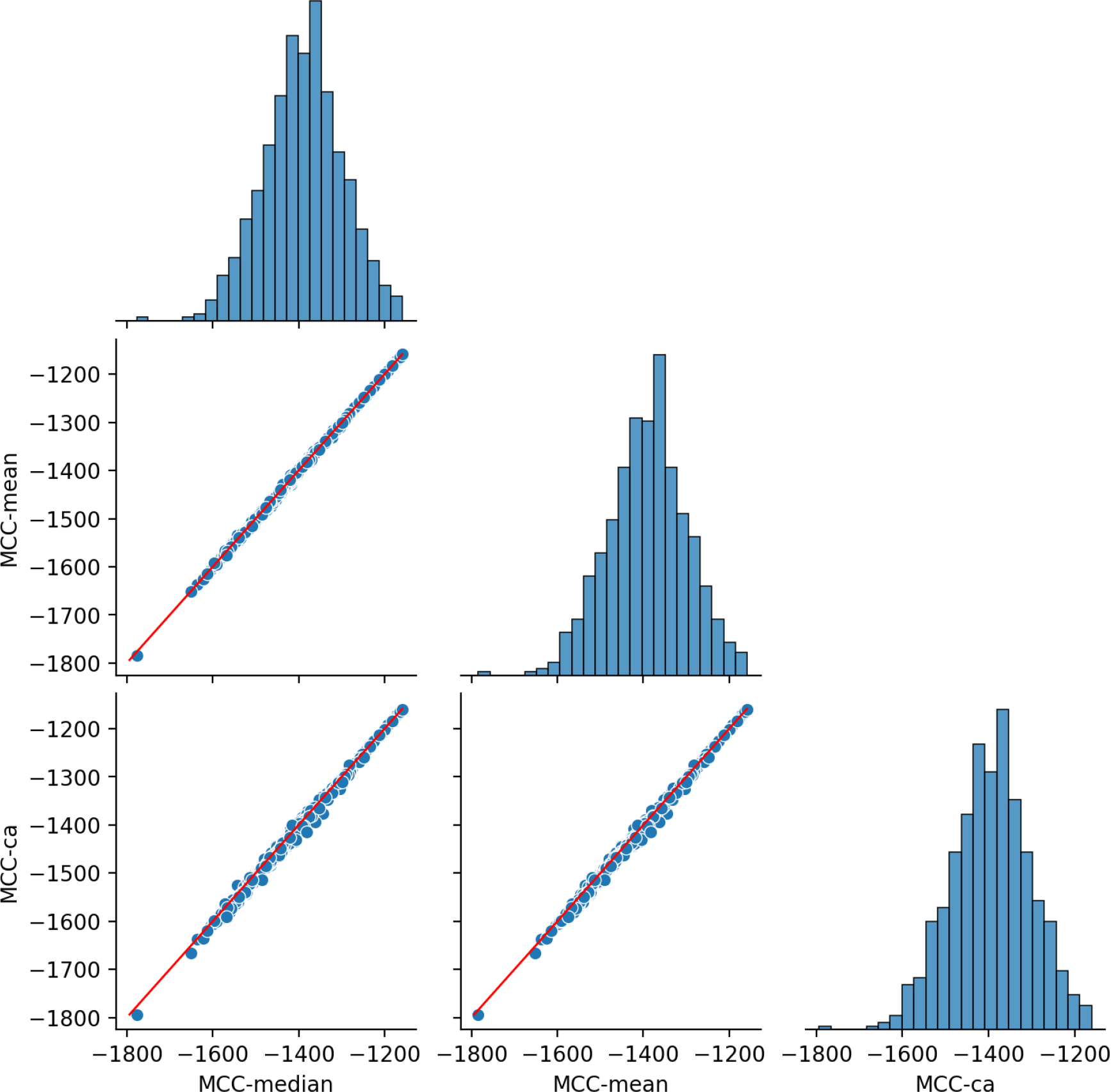
Comparing the log-likelihood values for different MCC branch length annotations. Deviation of the red diagonal line in the scatter plots indicate which tree annotation achieves higher log-likelihood values. The histograms show a distribution of the log-likelihood values for each of the annotations.

### 5.9. Detailed scatter plots for summary comparison plot

The main paper shows a summary of the following outcomes (Figure 16) of comparing the MCC and our centroid approximation in a histogram plot (Figure 3). Here we add the path difference metric [79] to the list of error measures. For it, the MCC always outperforms the centroid tree, however the values of the distance are orders of magnitude different especially for simulations with more taxa. We recognize that this is a strange behaviour of the metric and we are unaware as to why this is the case leaving us without a conclusion on this result.

**Figure 16.**
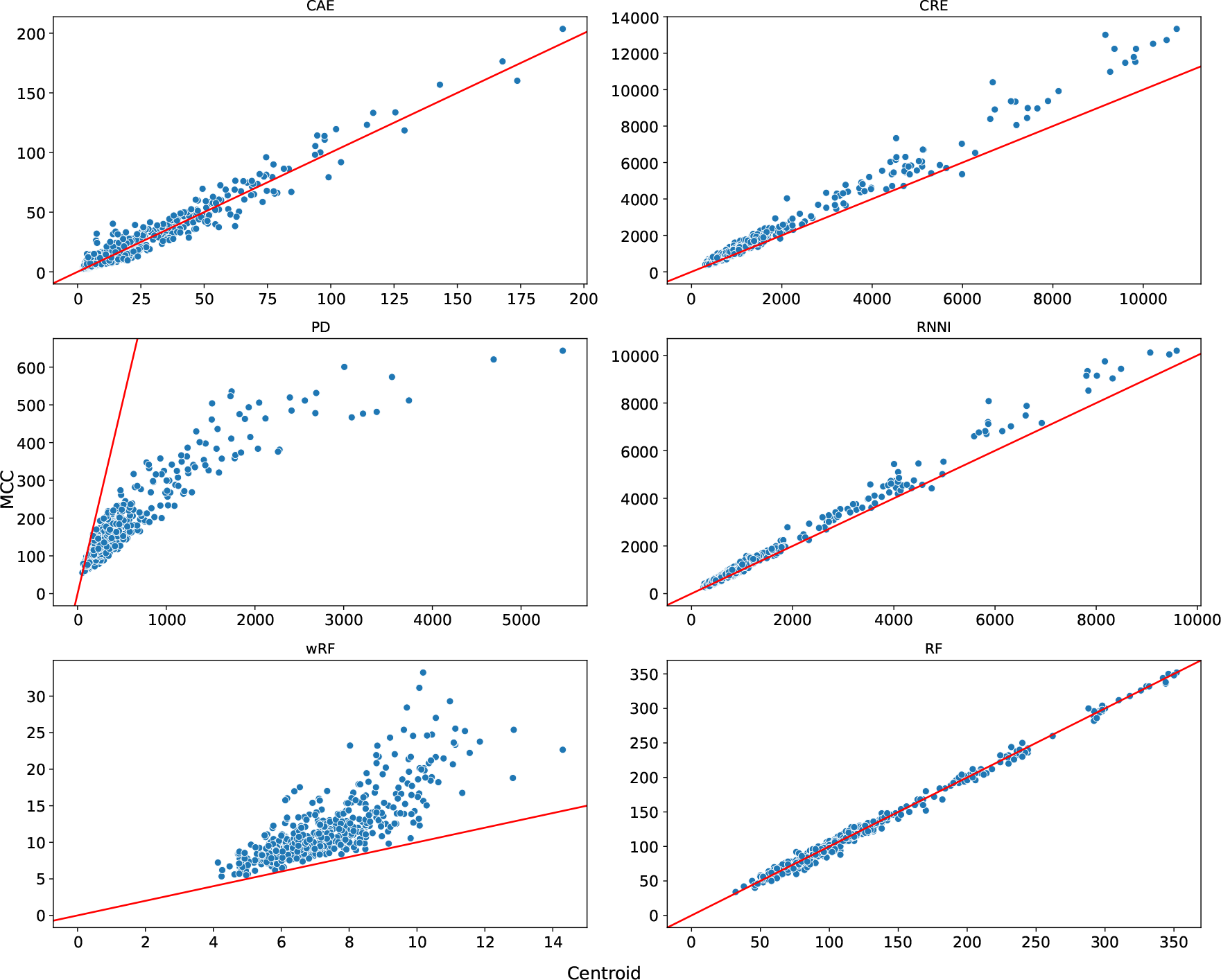
Comparison of the different error measures, the red diagonal is the identity line. The error measures displayed here are clade age error (CAE), clade rank error (CRE), path difference metric (PD) [79], RNNI metric, weighted Robinson Foulds metric (wRF) and the Robinson Foulds metric (RF).

### 5.10. More comparison plots

As stated in the main paper we only focused on simulations with more or equal to 50 taxa. The plot Figure 17 shows a comparison when datasets with fewer taxa are included. In Figure 18 a comparison using the median branch length annotation for the MCC tree can be seen. For that comparison the bars are divided by whether or not the MCC tree considered contains negative branch lengths or not. The plot shows that a vast majority of the MCC annotations contain negative lengths.

**Figure 17.**
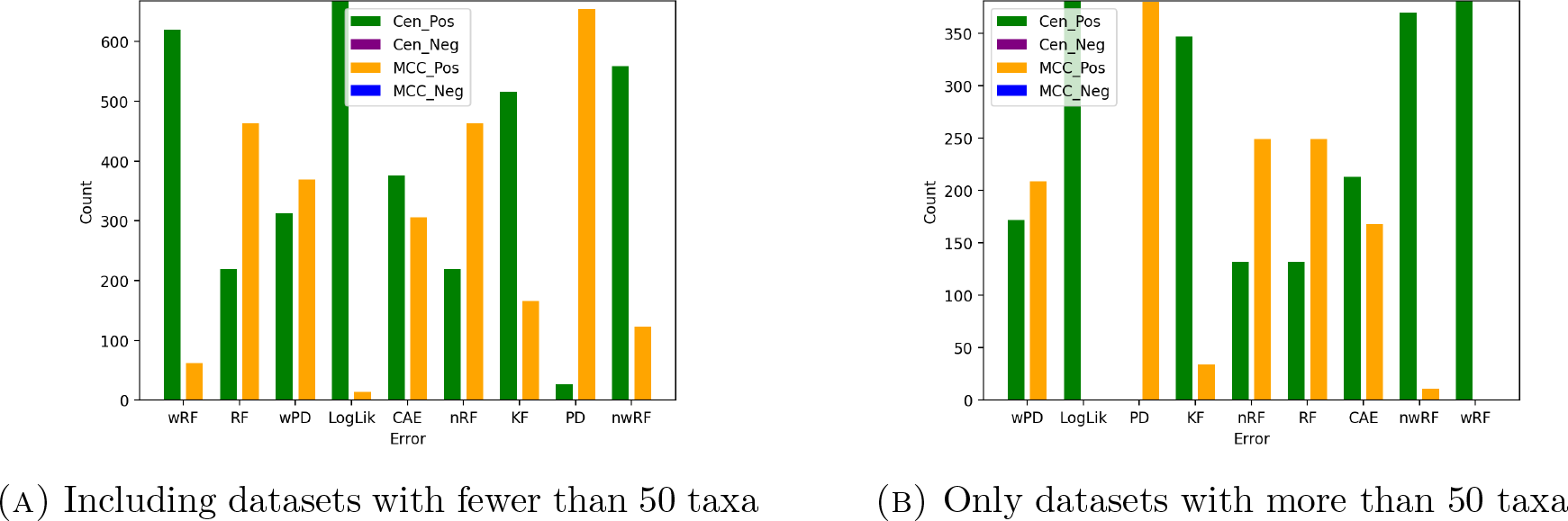
The error measure comparison slightly changes when small taxa datasets (fewer than 50) are included. This is because for these datasets it can often be the case, especially for data with fewer than 20 taxa, that the MCC tree and our centroid approximation are identical. The plot (A) includes simulations with 10, 12, 15, 20, 25, 30 and 35 taxa.

**Figure 18.**
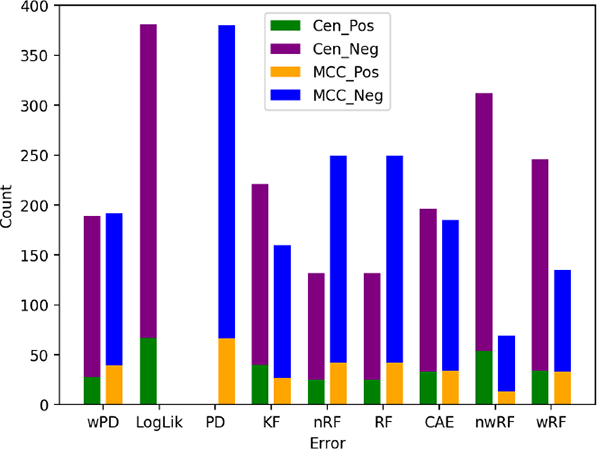
Comparing the error measures with the median branch length annotation for the MCC tree. Bars are split when the considered MCC tree has negative branch lengths, hence the identifier *Neg* (purple and blue bars) always implies negative branches in the MCC tree for the respective simulated dataset.

### 5.11. More Smoothness Plots

We analyse the smoothness property for the RNNI, BHV, Kendall-Colijn and Robinson-Foulds space by considering likelihood and posterior probabilities of a set of trees from following simulation: We simulate an alignment of 800bp for 100 taxa and run a BEAST2 analysis for 1, 000 iterations, logging every tree. Note that for the hill climbing samples we discard the first 500 samples as burn-in and for samples of the peak of the distribution we discard 500, 000 trees from the start of the chain. For these pairs of tree we then consider the relative distance (relative to largest observed distance) and the relative observed difference in the two log parameters resulting in the presented plot.

In Figure 19 we evaluate the smoothness of the RNNI and BHV treespaces at the tail end of a chain, i.e. while it is sampling trees from the peak of a distribution. Here the log-likelihood and posterior probability do not coincide like they do in the hill climbing example in the main paper. In Figure 20 the smoothness plot for both the hill climbing and the peak tree samples is presented for the Robinson-Foulds and the Kendall-Colijn distance.

**Figure 19.**
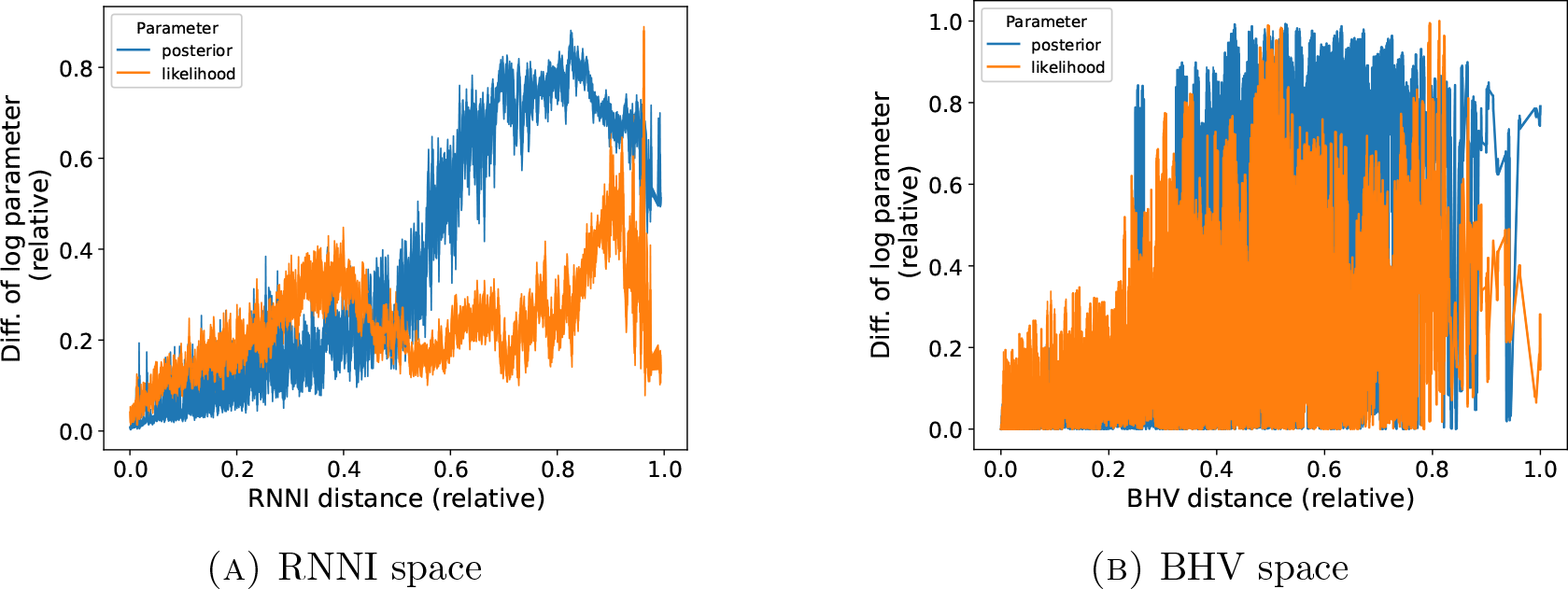
Smoothness assessment for samples from the peak of the distribution, i.e. samples from the end of a MCMC analysis.

**Figure 20.**
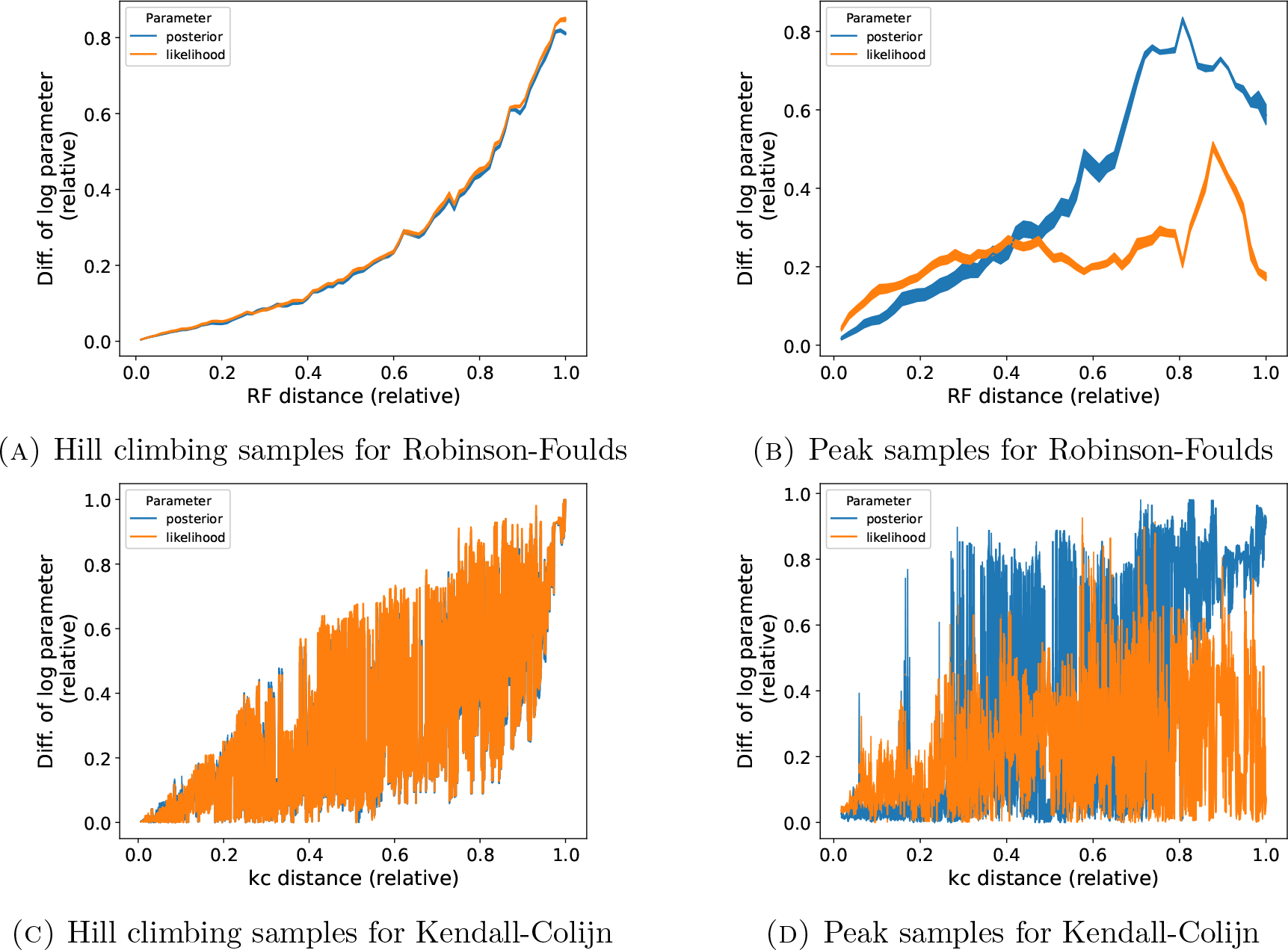
Smoothness assessment using the Robinson-Foulds and Kendall-Colijn distance.

### 5.12. Real data application

**Figure 21.**
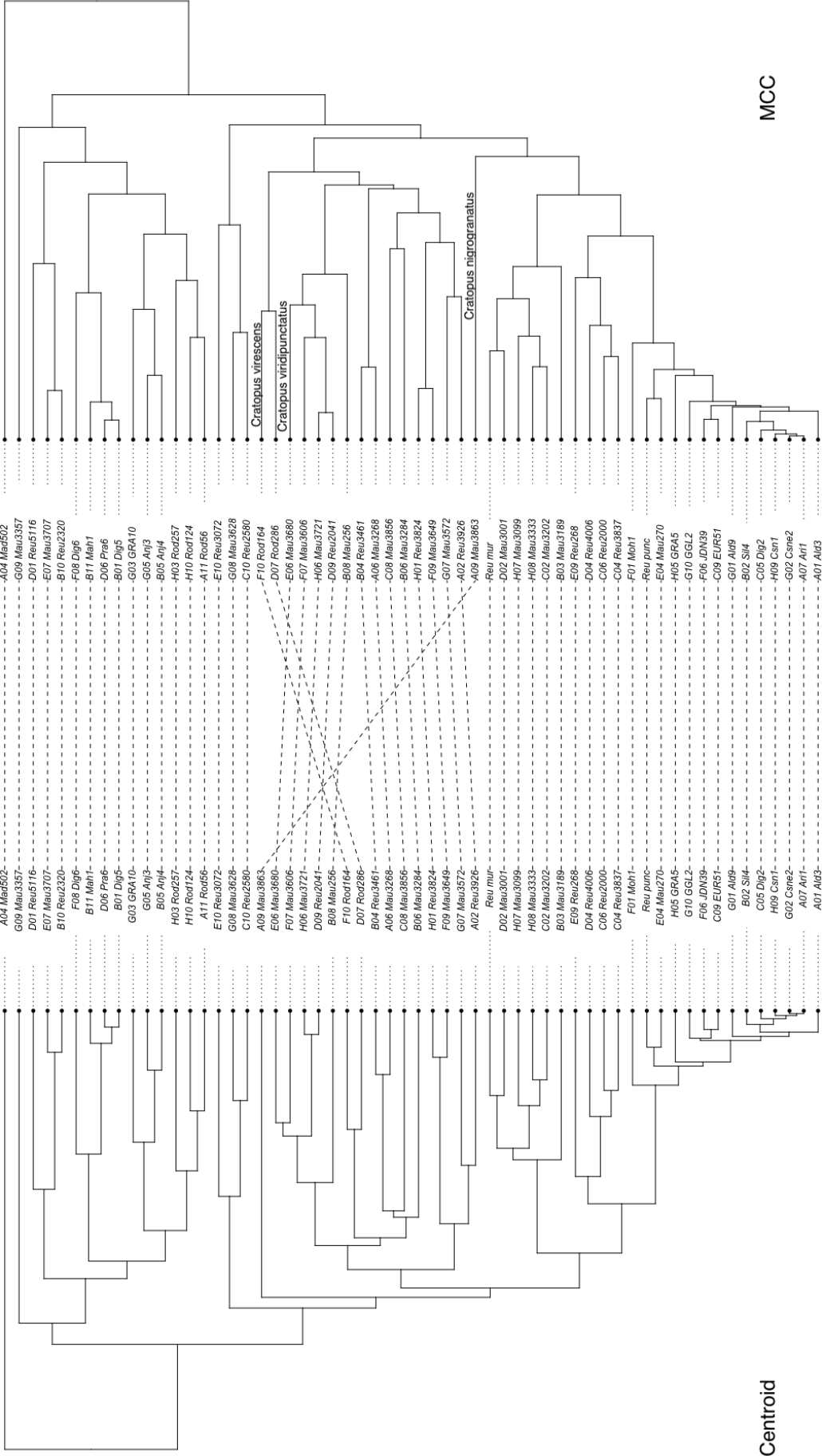
On this weevil dataset the trees differ in the position of three species as discussed in the main paper.

## Footnotes

1 For this result we use a minor modification to the Centroid algorithm that follows every possible path, i.e. every neighbour with lower SoS is pursued by the next iteration of the algorithm.

## References

1. Felsenstein, J. Inferring phylogenies (Sinauer associates Sunderland, MA, 2004).

2. Saitou, N. & Nei, M. The neighbor-joining method: a new method for reconstructing phylogenetic trees. Molecular biology and evolution 4, 406–425 (1987).

3. Fitch, W. M. Toward defining the course of evolution: minimum change for a specific tree topology. Systematic Biology 20, 406–416 (1971).

4. Swofford, D. *PAUP\**. Phylogenetic Analysis Using Parsimony (* and Other Methods). Version 4. 2003, *Sunderland, Massachusetts* 1999.

5. Felsenstein, J. Evolutionary trees from DNA sequences: a maximum likelihood approach. Journal of molecular evolution 17, 368–376 (1981).

6. Stamatakis, A. RAxML version 8: a tool for phylogenetic analysis and post-analysis of large phylogenies. Bioinformatics 30, 1312–1313 (2014).

7. Nguyen, L.-T., Schmidt, H. A., Von Haeseler, A. & Minh, B. Q. IQ-TREE: a fast and effective stochastic algorithm for estimating maximum-likelihood phylogenies. Molecular biology and evolution 32, 268–274 (2015).

8. Huelsenbeck, J. P. & Ronquist, F. MRBAYES: Bayesian inference of phylogenetic trees. Bioinformatics 17, 754–755 (2001).

9. Drummond, A. J. & Rambaut, A. BEAST: Bayesian evolutionary analysis by sampling trees. BMC Evolutionary Biology 7, 214. https://doi.org/10.1186/1471-2148-7-214 (2007).

10. A. Bouckaert, R., Vaughan, T. G., Barido-Sottani, J., Ducĥene, S., Fourment, M., Gavryushkina, A., Heled, J., Jones, G., Kühnert, D., De Maio, N., Matschiner, M., Mendes, F. K., Müller, N. F., Ogilvie, H. A., du Plessis, L., Popinga, A., Rambaut, A., Rasmussen, D., Siveroni, I., Suchard, M. A., Wu, C.-H., Xie, D., Zhang, C., Stadler, T. & Drummond, J. BEAST 2.5: An advanced software platform for Bayesian evolutionary analysis. eng. PLoS Comput Biol 15, e1006650. issn: 1553-7358 (Electronic); 1553-734X (Print); 1553-734X (Linking) (Apr. 2019).

11. Hohna, S., Heath, T. A., Boussau, B., Landis, M. J., Ronquist, F. & Huelsenbeck, J. P. Probabilistic graphical model representation in phylogenetics. Systematic biology 63, 753–771 (2014).

12. Maddison, D. R. The discovery and importance of multiple islands of most-parsimonious trees. Systematic Biology 40, 315–328 (1991).

13. Steel, M. The maximum likelihood point for a phylogenetic tree is not unique. Systematic Biology 43, 560–564 (1994).

14. Sanderson, M. J., McMahon, M. M. & Steel, M. Terraces in phylogenetic tree space. Science 333, 448–450 (2011).

15. Sanderson, M. J., McMahon, M. M., Stamatakis, A., Zwickl, D. J. & Steel, M. Impacts of terraces on phylogenetic inference. Systematic biology 64, 709–726 (2015).

16. Holmes, S. P. et al. Phylogenies: an overview. IMA Volumes in mathematics and its applications 112, 81–118 (1999).

17. Kim, J. Slicing hyperdimensional oranges: the geometry of phylogenetic estimation. Molecular phylogenetics and evolution 17, 58–75 (2000).

18. Billera, L. J., Holmes, S. P. & Vogtmann, K. Geometry of the Space of Phylogenetic Trees. Advances in Applied Mathematics 27, 733–767. issn: 0196-8858. https://www.sciencedirect.com/science/article/pii/S0196885801907596 (2001).

19. Moulton, V. & Steel, M. Peeling phylogenetic ‘oranges’. Advances in Applied Mathematics 33, 710–727 (2004).

20. Gill, J., Linusson, S., Moulton, V. & Steel, M. A regular decomposition of the edge-product space of phylogenetic trees. Advances in Applied Mathematics 41, 158–176 (2008).

21. Gavryushkin, A. & Drummond, A. J. The space of ultrametric phylogenetic trees. Journal of theoretical biology 403, 197–208 (2016).

22. Kendall, M. & Colijn, C. Mapping Phylogenetic Trees to Reveal Distinct Patterns of Evolution. en. Mol. Biol. Evol. 33, 2735–2743 (Oct. 2016).

23. Lin, B., Sturmfels, B., Tang, X. & Yoshida, R. Convexity in tree spaces. SIAM Journal on Discrete Mathematics 31, 2015–2038 (2017).

24. Gavryushkin, A., Whidden, C. & Matsen, F. A. The combinatorics of discrete time-trees: theory and open problems. Journal of Mathematical Biology 76, 1101–1121. https://doi.org/10.1007/s00285-017-1167-9 (2018).

25. Feragen, A. & Nye, T. in Riemannian Geometric Statistics in Medical Image Analysis 299–342 (Elsevier, 2020).

26. Collienne, L., Elmes, K., Fischer, M., Bryant, D. & Gavryushkin, A. Discrete coalescent trees. J Math Biol 83, 60. issn: 1432-1416 (Electronic) 0303-6812 (Linking).https://www.ncbi.nlm.nih.gov/pubmed/34739608 (2021).

27. Garba, M. K., Nye, T. M., Lueg, J. & Huckemann, S. F. Information geometry for phylogenetic trees. Journal of Mathematical Biology 82, 1–39 (2021).

28. Holmes, S. Statistics for phylogenetic trees. Theoretical Population Biology 63, 17–32. issn: 0040-5809. https://www.sciencedirect.com/science/article/pii/ S0040580902000059 (2003).

29. Nye, T. M. Principal components analysis in the space of phylogenetic trees. The Annals of Statistics, 2716–2739 (2011).

30. Baćak, M. Computing medians and means in Hadamard spaces. SIAM journal on optimization 24, 1542–1566 (2014).

31. Benner, P., Bacak, M. & Bourguignon, P.-Y. Point estimates in phylogenetic reconstructions. Bioinformatics 30, i534–i540 (2014).

32. Miller, E., Owen, M. & Provan, J. S. Polyhedral computational geometry for averaging metric phylogenetic trees. Advances in Applied Mathematics 68, 51–91. issn: 0196-8858. https://www.sciencedirect.com/science/article/pii/S0196885815000470 (2015).

33. Nye, T. M. Convergence of random walks to Brownian motion on cubical complexes. arXiv preprint arXiv:1508.02906 (2015).

34. Lin, B., Monod, A. & Yoshida, R. Tropical foundations for probability & statistics on phylogenetic tree space (2018).

35. Willis, A. & Bell, R. Uncertainty in phylogenetic tree estimates. Journal of Computational and Graphical Statistics 27, 542–552 (2018).

36. Willis, A. Confidence sets for phylogenetic trees. Journal of the American Statistical Association 114, 235–244 (2019).

37. Yoshida, R., Zhang, L. & Zhang, X. Tropical principal component analysis and its application to phylogenetics. Bulletin of mathematical biology 81, 568–597 (2019).

38. Page, R., Yoshida, R. & Zhang, L. Tropical principal component analysis on the space of phylogenetic trees. Bioinformatics 36, 4590–4598 (2020).

39. McMorris, F. R. & Steel, M. A. The complexity of the median procedure for binary trees in New Approaches in Classification and Data Analysis (eds Diday, E., Lechevallier, Y., Schader, M., Bertrand, P. & Burtschy, B.) (Springer Berlin Heidelberg), 136–140.isbn: 978-3-642-51175-2.

40. Heled, J. & Bouckaert, R. R. Looking for trees in the forest: summary tree from posterior samples. BMC Evol Biol 13, 221. issn: 1471-2148 (Electronic) 1471-2148 (Linking). https://www.ncbi.nlm.nih.gov/pubmed/24093883 (2013).

41. Hotz, T., Skwerer, S., Huckemann, S., Le, H., Marron, J. S., Mattingly, J. C., Miller, E., Nolen, J., Owen, M. & Patrangenaru, V. Sticky central limit theorems on open books. The Annals of Applied Probability 23, 2238–2258 (2013).

42. Matsen, F. A. A geometric approach to tree shape statistics. Systematic biology 55, 652–661 (2006).

43. Barthelemy, J.-P. & McMorris, F. R. The median procedure for n-trees. Journal of Classification 3, 329–334. issn: 0176-4268 (1986).

44. Kim, J., Rosenberg, N. A. & Palacios, J. A. Distance metrics for ranked evolutionary trees. Proceedings of the National Academy of Sciences 117, 28876–28886 (2020).

45. Owen, M. & Provan, J. S. A fast algorithm for computing geodesic distances in tree space. IEEE/ACM Trans Comput Biol Bioinform 8, 2–13. issn: 1557-9964 (Electronic) 1545-5963 (Linking). https://www.ncbi.nlm.nih.gov/pubmed/21071792 (2011).

46. Sturm, K.-T. Probability measures on metric spaces of nonpositive. Heat Kernels and Analysis on Manifolds, Graphs, and Metric Spaces: Lecture Notes from a Quarter Program on Heat Kernels, Random Walks, and Analysis on Manifolds and Graphs: April 16-July 13, 2002, Emile Borel Centre of the Henri Poincaŕe Institute, Paris, France 338, 357 (2003).

47. Barden, D., Le, H. & Owen, M. Central limit theorems for Fŕechet means in the space of phylogenetic trees. Electronic journal of probability 18, 1–25 (2013).

48. Lueg, J., Garba, M. K., Nye, T. M. W. & Huckemann, S. F. Wald Space for Phylogenetic Trees in Geometric Science of Information (eds Nielsen, F. & Barbaresco, F.) (Springer International Publishing), 710–717. isbn: 978-3-030-80209-7.

49. Bryant, D. A classification of consensus methods for phylogenetics. DIMACS series in discrete mathematics and theoretical computer science 61, 163–184 (2003).

50. Brown, D. G. & Owen, M. Mean and Variance of Phylogenetic Trees. Syst Biol 69, 139–154. issn: 1076-836X (Electronic) 1063-5157 (Linking). https://www.ncbi.nlmnih.gov/pubmed/31165169 (2020).

51. Rajanala, S. & Palacios, J. A. Statistical summaries of unlabelled evolutionary trees and ranked hierarchical clustering trees Electronic Article. June 2021. https://ui. adsabs.harvard.edu/abs/2021arXiv210602724R.

52. Robinson, D. F. & Foulds, L. R. in *Combinatorial mathematics VI* 119–126 (Springer, 1979).

53. Robinson, D. F. Comparison of labeled trees with valency three. Journal of combinatorial theory, Series B 11, 105–119 (1971).

54. Whidden, C. & Matsen 4th, F. A. Quantifying MCMC exploration of phylogenetic tree space. Syst. Biol. 64, 472–491 (May 2015).

55. Dasgupta, B., He, X., Jiang, T., Li, M. & Tromp, J. On Computing the Nearest Neighbor Interchange Distance. 55 (Sept. 2000).

56. Bordewich, M. & Semple, C. On the Computational Complexity of the Rooted Subtree Prune and Regraft Distance. Annals of Combinatorics 8, 409–423. https://doi.org/10.1007/s00026-004-0229-z (2005).

57. Bansal, M. S., Burleigh, J. G., Eulenstein, O. & Ferńandez-Baca, D. Robinson-foulds supertrees. Algorithms for molecular biology 5, 1–12 (2010).

58. Whidden, C., Beiko, R. G. & Zeh, N. Fixed-Parameter Algorithms for Maximum Agreement Forests. SIAM J. Comput. 42, 1431–1466 (Jan. 2013).

59. Smith, M. R. Information theoretic generalized Robinson–Foulds metrics for comparing phylogenetic trees. Bioinformatics 36, 5007–5013 (2020).

60. Smith, M. R. Robust analysis of phylogenetic tree space. Systematic Biology 71, 1255–1270 (2022).

61. Sokal, R. R. A statistical method for evaluating systematic relationships. *Univ. Kansas*, Sci. Bull. 38, 1409–1438 (1958).

62. Collienne, L. & Gavryushkin, A. Computing nearest neighbour interchange distances between ranked phylogenetic trees. Journal of Mathematical Biology 82, 1–19 (2021).

63. Summarizing posterior trees, BEAST2 https://www.beast2.org/summarizing-posterior-trees/.

64. Paradis, E. & Schliep, K. ape 5.0: an environment for modern phylogenetics and evolutionary analyses in R. Bioinformatics 35, 526–528 (2019).

65. Schliep, K. phangorn: phylogenetic analysis in R. Bioinformatics 27, 592–593. https://doi.org/10.1093/bioinformatics/btq706 (2011).

66. Jukes, T. H., Cantor, C. R. & Munro, H. N. in *Mammalian Protein Metabolism* 21–132 (Academic Press, 1969). isbn: 978-1-4832-3211-9. https://www.sciencedirect.com/science/article/pii/B9781483232119500097.

67. Rambaut, A., Drummond, A. J., Xie, D., Baele, G. & Suchard, M. A. Posterior Summarization in Bayesian Phylogenetics Using Tracer 1.7. Systematic Biology 67, 901–904. issn: 1063-5157. eprint: https://academic.oup.com/sysbio/article-pdf/67/5/901/25517397/syy032.pdf. (Apr. 2018).

68. Warren, D., Geneva, A. & Lanfear, R. RWTY (R We There Yet): An R package for examining convergence of Bayesian phylogenetic analyses R package version 1.0.2 (2017), 1016–1020. https://CRAN.R-project.org/package=rwty.

69. Bilderbeek, R. J. & Etienne, R. S. babette: BEAUti 2, BEAST 2 and Tracer for R. *Methods in Ecology and Evolution*. https://doi.org/10.1111/2041-210X.13032 (2018).

70. Mann, H. B. & Whitney, D. R. On a test of whether one of two random variables is stochastically larger than the other. The annals of mathematical statistics, 50–60 (1947).

71. Virtanen, P., Gommers, R., Oliphant, T. E., Haberland, M., Reddy, T., Cournapeau, D., Burovski, E., Peterson, P., Weckesser, W., Bright, J., van der Walt, S. J., Brett, M., Wilson, J., Millman, K. J., Mayorov, N., Nelson, A. R. J., Jones, E., Kern, R., Larson, E., Carey, C. J., Polat, İ., Feng, Y., Moore, E. W., VanderPlas, J., Laxalde, D., Perktold, J., Cimrman, R., Henriksen, I., Quintero, E. A., Harris, C. R., Archibald, A. M., Ribeiro, A. H., Pedregosa, F., van Mulbregt, P. & SciPy 1.0 Contributors. SciPy 1.0: Fundamental Algorithms for Scientific Computing in Python. Nature Methods 17, 261–272 (2020).

72. Robinson, D. F. & Foulds, L. R. Comparison of phylogenetic trees. Mathematical Biosciences 53, 131–147. issn: 0025-5564. https://www.sciencedirect.com/science/article/pii/0025556481900432 (1981).

73. Kolipakam, V., Jordan, F. M., Dunn, M., Greenhill, S. J., Bouckaert, R., Gray, R. D. & Verkerk, A. A Bayesian phylogenetic study of the Dravidian language family. Royal Society open science 5, 171504 (2018).

74. Kitson, J. J., Warren, B. H., Thebaud, C., Strasberg, D. & Emerson, B. C. Community assembly and diversification in a species-rich radiation of island weevils (Coleoptera: Cratopini). Journal of Biogeography 45, 2016–2026 (2018).

75. Alves, J. M., Prado-Lopez, S., Cameselle-Teijeiro, J. M. & Posada, D. Rapid evolution and biogeographic spread in a colorectal cancer. Nature communications 10, 5139 (2019).

76. Mooers, A. O. & Heard, S. B. Inferring Evolutionary Process from Phylogenetic Tree Shape. The Quarterly Review of Biology 72, 31–54. eprint: https://doi.org/10. 1086/419657. https://doi.org/10.1086/419657 (1997).

77. Schwarz, R. F., Ng, C. K., Cooke, S. L., Newman, S., Temple, J., Piskorz, A. M., Gale, D., Sayal, K., Murtaza, M., Baldwin, P. J., et al. Spatial and temporal heterogeneity in high-grade serous ovarian cancer: a phylogenetic analysis. PLoS medicine 12, e1001789 (2015).

78. Berling, L. Supplementary Centroid Code https://github.com/bioDS/Centroid-Code.

79. Steel, M. A. & Penny, D. Distributions of tree comparison metrics—some new results. Systematic biology 42, 126–141 (1993).

